# Trichome acylsugar defenses are equally effective against leaf-mining and free-living insects

**DOI:** 10.64898/2026.05.27.728314

**Authors:** Zongyan Sun, Xiaochan Pan, Simin Bi, Yirui Zhang, Junling Dun, Dina Mahesati, Yibin Lin, Ying Fu, Yang Gao, Xiaoyan Xu, Na Chen, Jiarui Xing, Wenfeng Ye, Minbiao Ji, Ian T. Baldwin, Jiancai Li

## Abstract

Trichomes are epidermal projections, crucial components of plant defense systems. Whether these epidermal defenses are effective against leafminers, which infest the internal mesophyll layers, remains unclear. Using the devasting invasive tomato pest, *Phthorimaea absoluta* (tomato leafminer), and the trichome-rich wild tomato, *Solanum pennellii*, we demonstrate that trichome produced acylsugars effectively inhibit the growth of endophagous *P. absoluta* larvae feeding inside leaves. High spatial resolution leaf- and frass-metabolite analysis by MALDI-MSI and Stimulated Raman Scattering microscopy revealed that trichome-synthesized acylsugars are translocated into the mesophyll tissues when larvae infest leaves. We identified an ATP-binding cassette (ABC) transporter, SpABCB5, as the mediator of this translocation and as an essential component of defense against leafminers. These data reveal that epidermal trichomes not only defend against free-living (ectophagous) herbivores through direct contact or rapid deposition of chemical defenses onto leaf surfaces, but also defend against endophagous leafminers by rapidly depositing chemical defenses into internal tissues. By delivering mines against leaf miners, trichomes offer new avenues for breeding crops with resistance to leafminers.

---

Leaf-mining insects represent a specialized guild of herbivores whose larvae feed and develop within the mesophyll, residing between the upper and lower epidermal layers throughout their immature stages [1]. This group encompasses economically devastating pests, such as leafminer moths (Lepidoptera) and leafminer flies (Diptera), which cause substantial damage to agricultural crops globally [2]. The endophytic lifestyle confers distinct ecological advantages: by remaining concealed within leaf tissues, leafminers effectively escape predation and parasitism, buffer against adverse environmental conditions, and, critically for agriculture, avoid direct exposure to contact insecticides [3]. This protected feeding niche severely compromises the efficacy of conventional chemical control measures, leading to substantial yield losses in crop production and environmental pollutions by overuse of pesticides [4]. Although enhancing plant resistance offers a theoretically robust strategy to counteract the inherent survival advantages of leafminers and reduce reliance on pesticides, the molecular mechanisms governing plant-leafminer interactions remain disproportionately understudied compared to those involving chewing or piercing-sucking herbivores [5].

The leaf epidermis serves as the primary interface and initial barrier against herbivory. Glandular trichomes, specialized epidermal protrusions prevalent across numerous plant families, function as dynamic “metabolic factories” dedicated to the biosynthesis and secretion of defensive specialized metabolites [6, 7]. These structures orchestrate a dual defensive strategy: physically impede insect locomotion and oviposition, and chemically assaulting herbivores through the release of volatile organic compounds that modulate insect behavior or recruit natural enemies, and deposition of non-volatile metabolites on the leaf surface [8]. While the role of these exudates in deterring surface-feeding insects is well-documented [9], a critical knowledge gap persists regarding their effectiveness against leaf-mining insects. Specifically, it remains unknown whether metabolites synthesized and secreted at the trichome apex can exert toxic or deterrent effects on larvae that rapidly transition from the leaf surface to the internal tissue, or if the mining behavior effectively circumvents this chemical arsenal.

Acylsugars, comprising a sugar core esterified with fatty acid chains, represent one of the most potent anti-herbivore metabolites produced by glandular trichomes in the Solanaceae family [10, 11]. Their biosynthesis is mediated by four consecutive BAHD acyltransferases (ASAT1-4), which catalyze the sequential attachment of fatty acid chains to the sugar core via ester bonds [12-14]. *Nicotiana benthamiana* plants harboring a mutation in the first dedicated acylsugar biosynthesis gene, *Nbasat1*, exhibit heightened susceptibility to whiteflies (*Bemisia tabaci*), aphids (*Macrosiphum euphorbiae*) and lepidopteran larvae *Helicoverpa zea* [15], all major pests of tomato production. The tomato leafminer, *Phthorimaea absoluta* (Meyrick), has recently emerged as a devastating global threat to tomato production, causing severe yield losses [16, 17]. Although previous studies have correlated resistance to *P. absoluta* with glandular trichome density and acylsugar content [18-20], it remains unclear whether acylsugars act as the direct causal agents of this resistance. Mechanistically, it is unknown how surface-localized acylsugars interact with a pest that spends the majority of its life cycle hidden within the leaf parenchyma.

In this study, we exploit the interaction between *Solanum pennellii*, a wild tomato species known for its high accumulation of acylsugars, and the tomato leafminer to dissect the interaction between trichome-derived metabolites and endophytic herbivores. Through the integration of genetic mutant and insect frass metabolite analysis, we establish a definitive causal link between acylsugars and resistance to tomato leafminer. Furthermore, high-resolution spatial metabolomics reveals a novel defense mechanism: insect feeding triggers the active transport of acyl sugars from trichomes into the mesophyll cells subtending trichomes, a process mediated by a specific ABC transporter. Mutants deficient in this transporter exhibit markedly impaired acylsugar translocation and compromised pest resistance. Collectively, these findings reveal that endophagous leafminers remain within the effective “firing range” of the trichome produced chemical defenses, that trichomes mine leaves in response to leafminer attack, and thereby provide new genetic targets for controlling leaf-mining pests.

## Result

### Trichome-derived acylsugars confer resistance against tomato leafminer

To elucidate the interaction of the endophytic tomato leafminer (*Phthorimaea absoluta*) and trichome-derived metabolites, we meticulously observed the larval feeding behavior on both the cultivated tomato variety (M82) and the wild tomato *Solanum pennellii* (LA0716), which is known for its high acylsugar accumulations. Our observations revealed that larval feeding does not directly contact either glandular or non-glandular trichomes (Fig. 1A and Movie 1). While M82 harbors substantial densities of type-V trichomes, LA0716 plants possess almost exclusively type-I and type-IV glandular trichomes, which are characterized by visible acylsugar droplets at their tips (Fig. 1B and S1A). Consistent with the higher density of acylsugar-producing trichomes (Fig. 1C), liquid chromatography-mass spectrometry (LC-MS) analysis confirmed that LA0716 accumulates significantly higher levels of acylsugars compared to M82 (Fig. 1D). Importantly, tomato leafminer larvae reared on wild tomato LA0716 gained only about 30% of body mass of those reared on M82 plants (Fig. 1E and S1B).

**Fig. 1.**
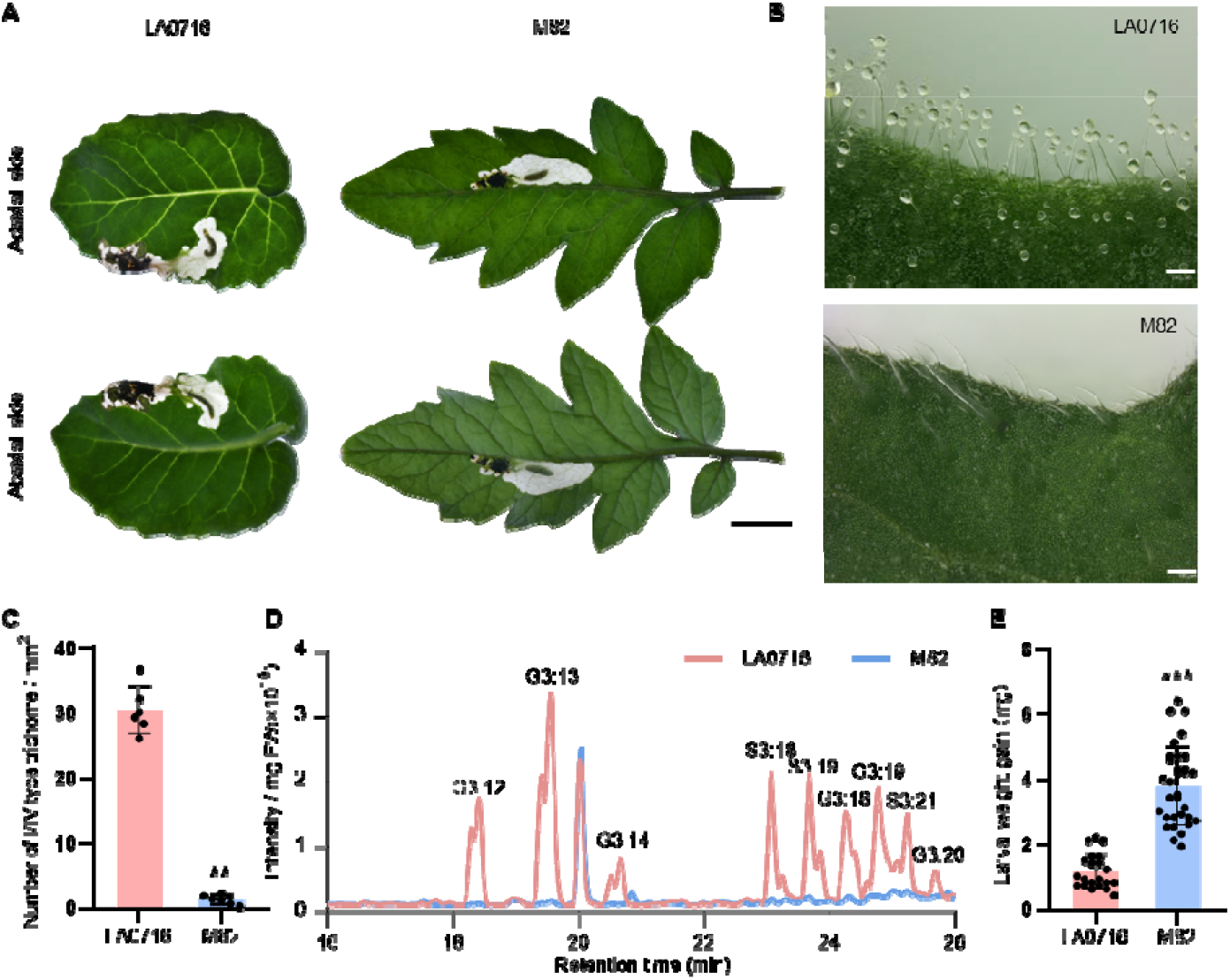
Acylsugar enriched *S. pennellii* is a poor host for leafminers. (A) Leaf damage caused by tomato leafminer on wild tomato *S. pennellii* LA0716 (left) and cultivated tomato M82 (right). Scale bar, 1 cm. (B) Macroscopic images of leaf epidermal surfaces from LA0716 (top) and M82 (bottom). Prominent acylsugar droplets are visible on the LA0716 surface. Scale bar, 100 μm. (C) Liquid chromatograph-mass spectrometry (LC-MS) chromatograms of leaf metabolite extracts from LA0716 and M82. Peaks corresponding to acylsugars are labeled. Acylsugars are named according to their sugar core (S, sucrose; G, glucose), number of acyl chains, and total acyl chain carbon count. Nomenclature rules are provided in Fig. S2 legend. (D) Density of type I and type IV trichomes (number per mm^2^; mean ± SE, *n* = 6) on leaves of LA0716 and M82. (E) Larval mass of *P. absoluta* (mean ± SE, *n* = 21 and 32, respectively) after 10 days of feeding on indicated plant genotypes. Statistical significance for C and E was determined using independent-sample Student-*t* tests (*** *P* < 0.01; *** *P* < 0.001).

To determine whether high acylsugar levels were responsible for the superior resistance of wild tomato, we generated knockout mutants of *SpASAT1*, the first dedicated gene in the acylsugar biosynthetic pathway (Fig. S2A), in wild tomato LA0716 background using CRISPR/Cas9 with two distinct sgRNAs. We obtained two independent T-DNA-free lines: one (*asat1#1*) containing a premature stop codon, and the other (*asat1#2*) harboring two fragment deletions, including one within the conserved HXXXD motif characteristic of BAHD acyltransferases (Fig. 2A and Fig. S2) [21]. The visible acylsugar droplets present on wild-type LA0716 trichomes were absent in both independent *asat1* mutant lines (Fig. 2B and S1C). Although the *ASAT1*-knockout did not significantly alter trichome density (Fig. S1D), it reduced total leaf acylsugar accumulations to approximately 0.2% of wild-type levels (Fig. 2C and 2D). Tomato leafminer larvae feeding on the *asat1* mutants gained about threefold more mass, compared with those feeding on wild-type LA0716 (Fig. 2E). Collectively, these data demonstrate that trichome-synthesized acylsugars are indeed causal factors in resistance against the endophagous tomato leafminer.

**Fig. 2.**
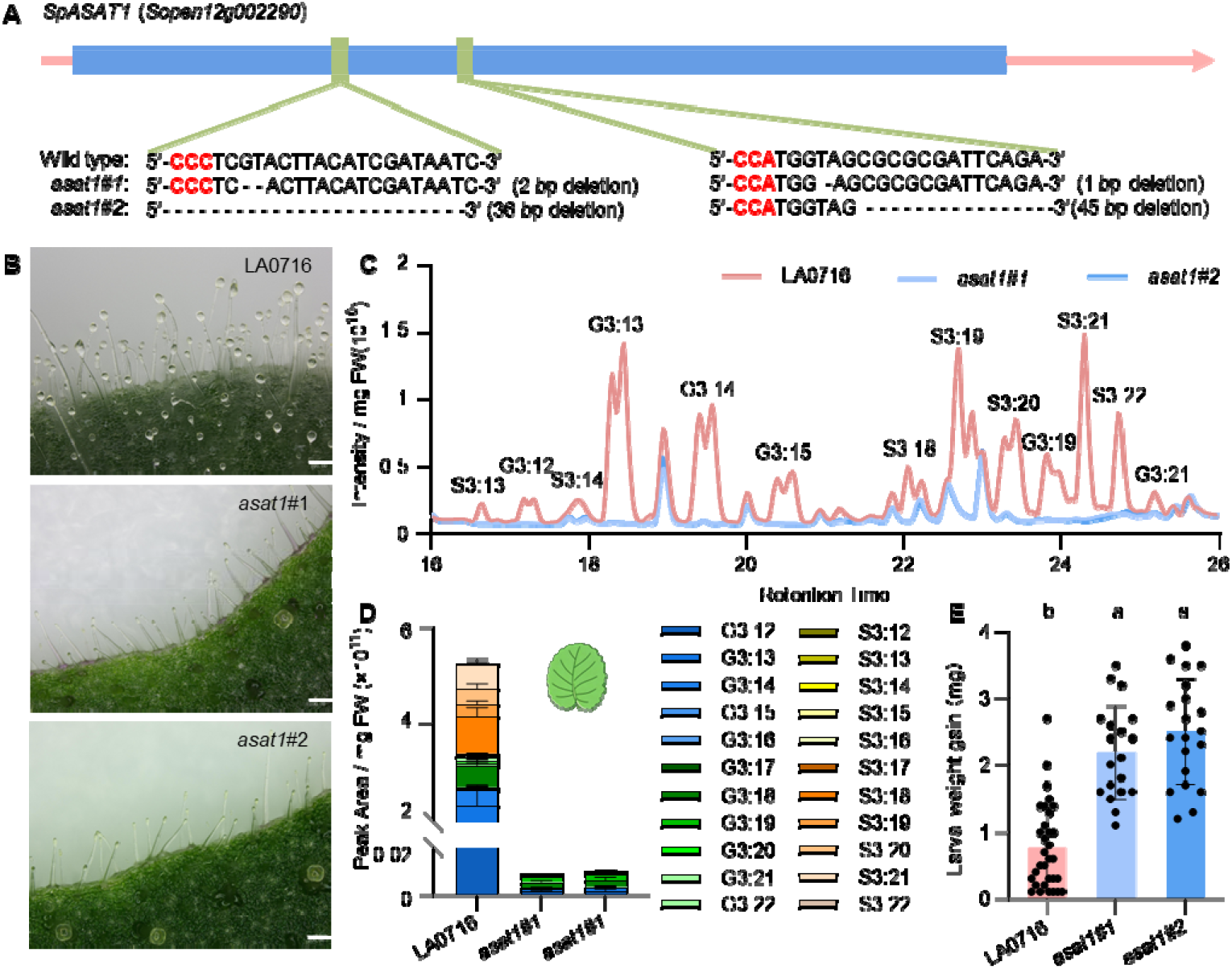
*SpASAT1* knockout dramatically reduces wild tomato LA0716 resistance to leafminers. (A) Schematic of *SpASAT1* gene mutation generation using CRISPR-Cas9-based genome editing. The two sgRNA target sequences and resulting mutation types are listed. Protospacer adjacent motif (PAM) sequences are highlighted in red. (B) Macroscopic images of leaf epidermal surfaces from LA0716 and two independent *asat1* mutant lines. Scale bar, 100 μm. (C) LC-MS chromatograms of leaf metabolite extracts from LA0716 and two independent *asat1* mutant lines. Peaks corresponding to acylsugars are labeled. (D) Relative abundance of acylsugars (mean + SE, *n* = 3) in the leaf extracts of the indicated genotypes. (E) Larval mass of *P. absoluta* (mean ± SE, *n* = 32, 20, and 21, respectively) after feeding on the indicated genotypes for 10 days. Different letters indicate significant differences (*P* < 0.05, one-way ANOVA with Tukey’s *post hoc* test).

### Trichome-synthesized acylsugars are mesophyll-translocated in response to leafminer attack

To investigate whether acylsugar are directly toxic to *P. absoluta*, we analyzed the metabolite composition of larval frass. To prevent contamination from the ubiquitous plant-surface acylsugars, larvae were dissected from their mines and transferred from host plants to clean vials, and only freshly produced frass was collected for metabolite analysis.

Substantial amounts of acylsugars were detected in the frass of larvae fed on LA0716 plants (Fig. 3A and 3B). Consistent with the near-absence of acylsugars in *asat1* mutant plants (Fig. 2C), frass from larvae feeding on these *asat1* mutant plants contained acylsugars at concentrations only 5% of that in the frass of larvae feeding on wild-type LA0716 plants (Fig. S4A). Although less than in the frass of larvae fed wild tomato LA0716 plants, acylsugars were also detected in the frass from cultivated M82 tomato fed larvae (Fig. S4B). These findings confirm that leafminers indeed ingest acylsugars despite their leaf-mining feeding habit.

**Fig. 3.**
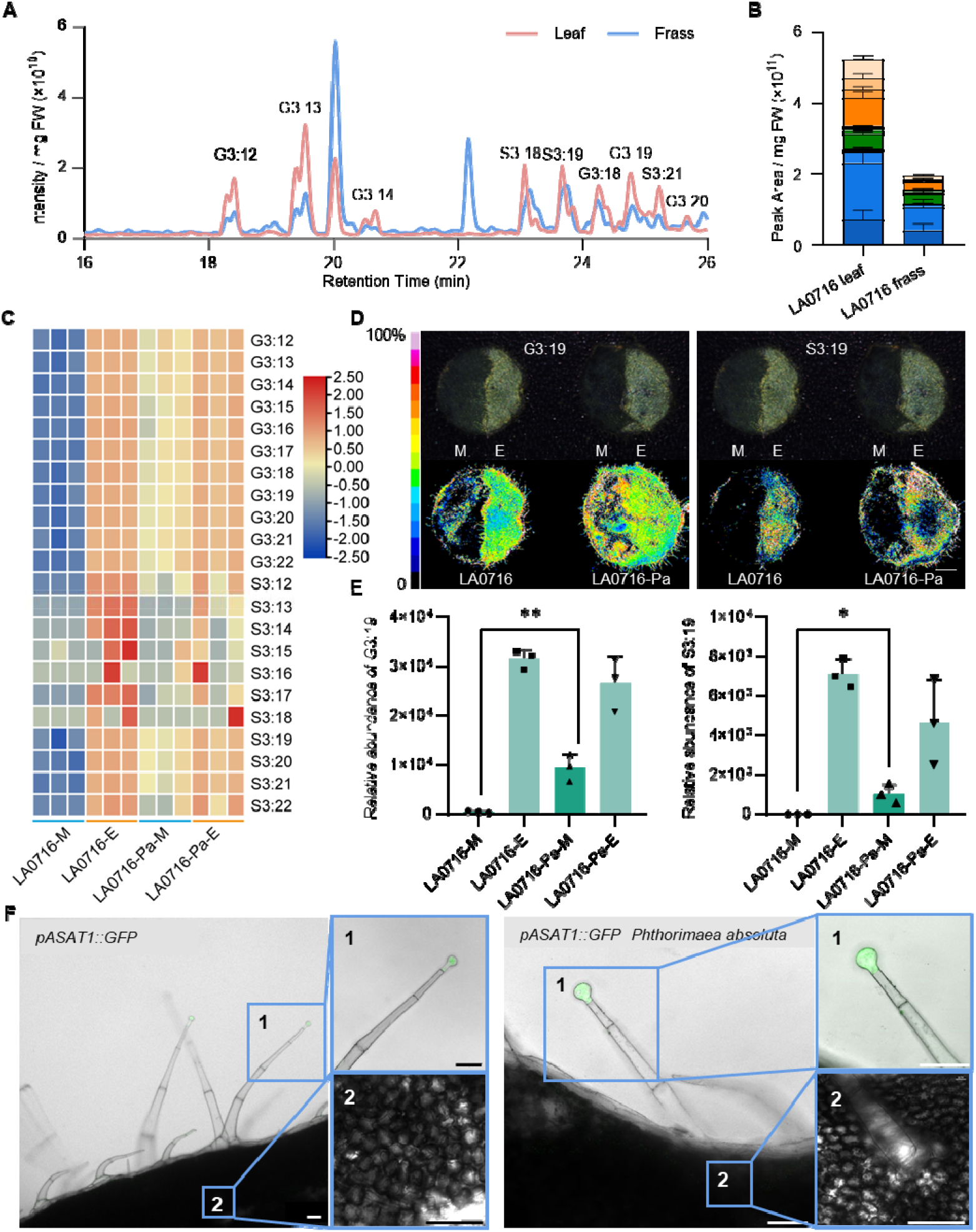
Acylsugars are translocated to mesophyll tissues in response to leafminer infestation. (A) LC-MS chromatograms of metabolite extracts from LA0716 leaves and larval frass collected from larvae fed on LA0716 plants. Peaks corresponding to acylsugars are labeled. (B) Relative abundance of acylsugars (mean + SE, *n* = 3) in extracts of LA0716 leaves and frass from larvae fed on LA0716. (C) Heatmap of signal intensities of acylsugars (mean, *n* = 3) detected by MALDI-MSI in mesophyll of uninfested LA0716 (LA0716-M), epidermis of uninfested LA0716 (LA0716-E), mesophyll of *P. absoluta*-infested LA0716 (LA0716-Pa-M), and epidermis of *P. absoluta*-infested LA0716 (LA0716-Pa-E). (D) Optical images of leaf discs used for MALDI-MSI analysis (top) and MALDI-MSI of representative acylsugars G3:19 (left) and S3:19 (right). The intensity of spectra representing each metabolite is visualized in false color. M and E indicate samples were harvested from mesophyll and epidermis, respectively. Scale bar, 2 mm. (E) Relative intensity of MALDI-MSI signals (mean + SE, *n* = 3) for representative acylsugars G3:19 (left) and S3:19 (right) from indicated treatments. (F) Confocal image showing GFP fluorescence driven by the *SlASAT1* promoter under non-infested (left) or *P. absoluta*-infested (right) conditions. Scale bar, 50 μm. Statistical significance for E was determined using independent-sample Student-*t* tests (* *P* < 0.05; ** *P* < 0.01).

Since *P. absoluta* larvae feed exclusively on mesophyll tissue after penetrating the epidermis as neonates, the presence of acylsugars in the frass of the fourth-instar larvae used for the experiments, implies that trichome-synthesized compounds are translocated from the trichome apices into the underlying mesophyll tissues. However, physically isolating mesophyll tissue without contamination from epidermal acylsugars is technically challenging. To overcome this, we employed matrix-assisted laser desorption/ionization-mass spectrometry imaging (MALDI-MSI) to map acylsugar distribution in leaf discs, comparing intact leaves with those in which the epidermis had been removed. The high signal intensities of various acylsugars from the intact epidermis confirmed the reliability of our MALDI-MSI protocol for acylsugar detection, including both glucose-core and sucrose-core acylsugars (Fig. 3C). Substantial acylsugar signals were detected in the mesophyll of leafminer-infested leaves, whereas non-infested leaves showed negligible mesophyll accumulation (Fig. 3D, 3E and S4).

To determine whether the acylsugars found in infested mesophyll tissues were translocated from trichomes or locally synthesized *de novo* in response to infestation, we monitored the spatial expression of *ASAT1*. A transgenic tomato plant expressing *GFP* under the control of the *ASAT1* promoter (*pASAT1::GFP*) in the M82 background [12] was used for expression monitoring. Consistent with previous research [12], GFP fluorescence was observed exclusively in the apical cell of glandular trichomes (Fig. 3F). Crucially, no expression was detected in mesophyll cells, regardless of leafminer infestation status. Collectively, these results demonstrate that trichome-derived acylsugars are actively translocated from trichome apices into mesophyll tissues in response to leafminer infestation.

### SpABCB5 mediates trichome-mesophyll acylsugar translocation

Given that the transporter expression is often co-regulated with cargo biosynthetic genes [22], we performed RNA-seq on seven distinct tissues of the wild tomato LA0716 (leaf, stem, root, flower, fruit, leaf trichome and stem trichome) to identify transporters responsible for acylsugar translocation from trichome apieces to the mesophyll tissues. K-means clustering (k = 20) of all detected genes identified a “trichome-specific” cluster containing 1703 genes (Fig. S6). Within this cluster, we identified seven ABC transporters (Fig. 4A). We selected the three most highly expressed, trichome-specific candidates, *SpABCB5, SpABCG6* and *SpABCG54*, for functional characterization by virus-induced gene silencing (VIGS). The success of VIGS procedure was confirmed by silencing the phytoene desaturase (PDS) gene, which resulted in a distinct bleaching phenotype (Fig S7A). LC-MS analysis of acylsugars in the leaves of these VIGS plants showed that the total acylsugar levels were reduced in *SpABCB5*-silenced plants, but not in plants silenced for the other two *ABCG* genes (Fig. S7B). Furthermore, bioassays with the tomato leafminer on these VIGS plants revealed that only silencing *SpABCB5* significantly compromised larval resistance in LA0716 plants (Fig. S7C). Additionally, among the four trichome highly expressed ABC transporters analyzed by qRT-PCR, only *SpABCB5* expression was induced by *P. absoluta* infestation (Fig. 4B and S7C). Collectively, these data indicate that *SpABCB5* is the most likely candidate responsible for acylsugar translocation from trichome to mesophyll.

**Fig. 4.**
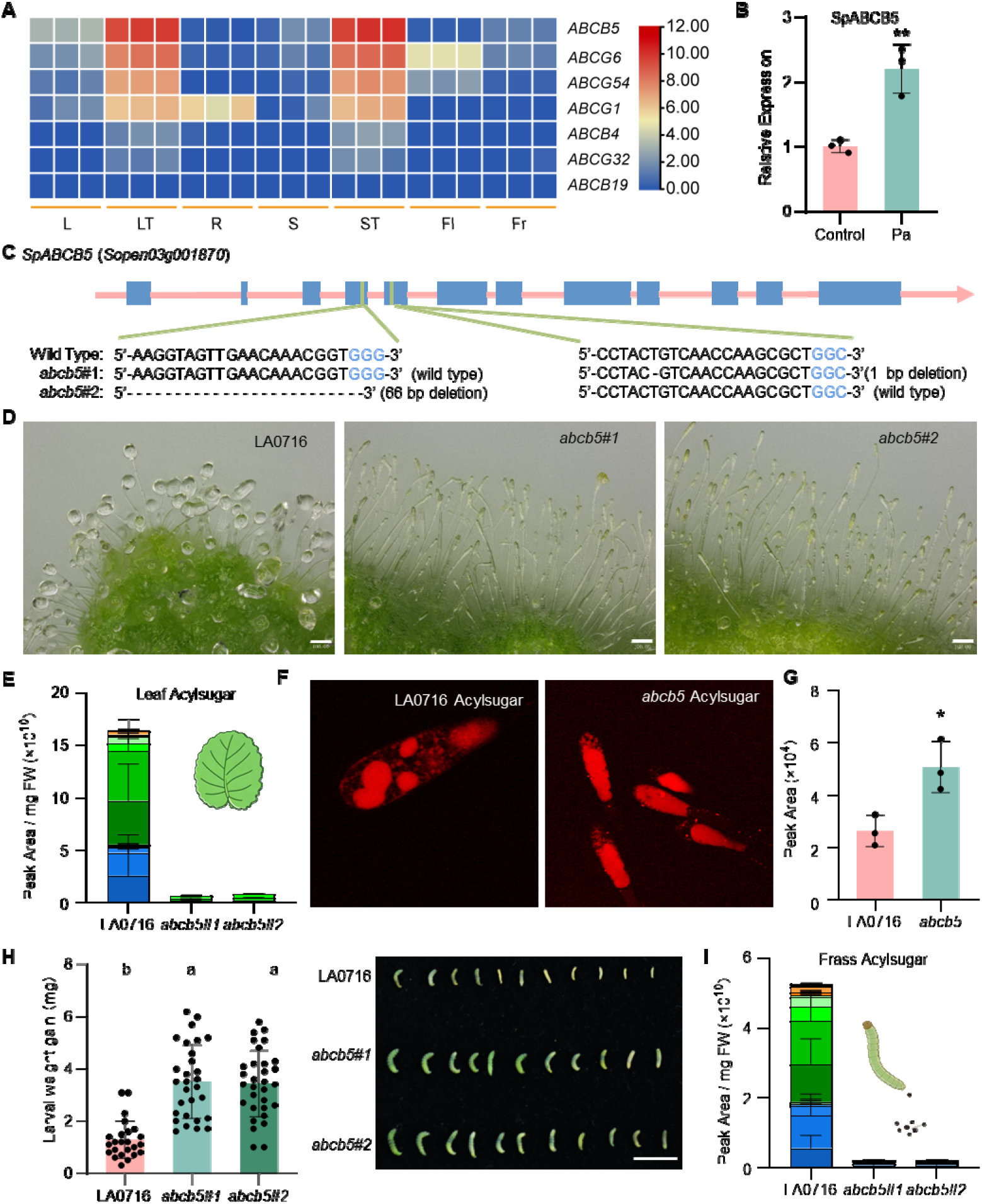
SpABCB5 contributes to the trichome-mesophyll acylsugar translocation. (A) Tissue-specific expression profiles of ABC transporters in “trichome-specific” cluster; heatmap coloring depicts the transcripts per million (TPM). L, leaf; LT, leaf trichome; R, root; S, stem; ST, stem trichome; Fl, flower; Fr, fruit. (B) The relative expression of SpABCB5 in the leaves of uninfested (control) or *P. absoluta*-infested (Pa) LA0716. (C) Schematic of *SpABCB5* gene mutation generation using CRISPR-Cas9-based genome editing. The two sgRNA target sequences and resulting mutation types are listed. PAM sequences are highlighted in blue. (D) Macroscopic images of leaf epidermal surfaces from LA0716 and two independent *abcb5* mutant lines. Scale bar, 100 μm. (E) Relative abundance of acylsugars (mean + SE, *n* = 3) in the leaf extracts of the indicated genotypes. (F) SRS image of trichome apical cell on wild-type LA0716 and abcb5 mutant lines at the acylsugar specific wavelengths. (G) Relative intensities of acylsugar signals (mean + SE, *n* = 3) in wild-type LA0716 and abcb5 mutant, harvested by SRS microscope. (H) Larval mass (mean ± SE, *n* = 24, 29, and 31, respectively) and representative image of *P. absoluta* after feeding on indicated genotypes for 10 days. Scale bar, 1 cm. (I) Relative abundance of acylsugars (mean + SE, *n* = 3) in extracts of frass from larvae fed on LA0716, and frass from larvae fed on the *abcb5* mutant. Statistical significance for B and G was determined using independent-sample student-*t* tests (\**P* < 0.05; \*\**P* < 0.01). Different letters indicate significant differences (*P* < 0.05, one-way ANOVA with Tukey’ s *post hoc* test).

To definitively examine the role of SpABCB5 in acylsugar translocation, we generated *SpABCB5* knockout mutants using CRISPR/Cas9 with two sgRNA targeting separate exons (Fig. 4C). We obtained two independent T-DNA-free lines: *abcb5#1*, harboring a 1-bp deletion causing a premature stop codon, and *abcb5#2*, containing a 66-bp deletion within a transmembrane domain (Fig. 4C and S8). The visible acylsugar droplets were absent on both mutant lines (Fig. 4D). LC-MS analysis confirmed that total leaf acylsugar content in *abcb5* mutant was reduced to approximately 5% of that of wild-type LA0716 leaves (Fig. 4E), despite no significant alteration in the densities of acylsugar-producing trichome (Fig. S9A).

To distinguish whether this dramatic reduction of acylsugars in *abcb5* mutants resulted from secretion blockage or direct impairment of biosynthesis, quantification of intracellular acylsugars is required. However, the longitudinal cross-sectional area of trichome apical cell (< 100 μm^2^) is below the resolution limit of current MALDI-MSI techniques. Leveraging the high sensitivity of stimulated Raman scattering (SRS) microscopy [23], we developed an SRS-based assay targeting distinct acylsugar vibrational signals at 1572 and 1615 cm^−1^ (Fig. S10). When tuned to these wavelengths, SRS microscopy revealed that acylsugar signals clustered into droplets that colocalized with vacuoles in brightfield images (Fig. 4F and S9B). While wild-type trichome apical cells typically contained multiple small acylsugar-filled vacuoles, *abcb5* mutant cells exhibited a single, enlarged vacuole with significantly higher SRS signal intensity (Fig. 4F, S9C and S9D). Quantitative SRS analysis demonstrated that trichome apical cells in *abcb5* mutant lines accumulated 94% more acylsugars than those in wild-type LA0716 plants (Fig. 4G). This higher intracellular accumulation, coupled with the drastic reduction in total leaf acylsugars, suggests that the blocked secretion caused by *abcb5* mutation triggers a feedback suppression of the biosynthetic pathway. Consistent with this hypothesis, the transcriptome analysis of *abcb5* mutant trichomes revealed that nearly all previously identified acylsugar metabolism genes [24] were downregulated, including those involved in the branch-chain amino acid (BCAA) pathway, fatty acid synthesis (FAS), acyl-activating enzymes (AAEs), ASATs, acylsugar hydrolases (ASHs), transporters, and transcriptional factors (Fig. S11A). Furthermore, enrichment analysis indicated that these differently expressed genes were significantly enriched in these pathways (Fig. S11B). These data collectively demonstrate that SpABCB5 facilitates acylsugar efflux from the cytosol to the apoplast.

To further elucidate the role of SpABCB5 in basipetal translocation into mesophyll tissues, we performed larval growth bioassays comparing *abcb5* mutants and wild-type LA0716. *P. absoluta* larvae feeding on *abcb5* mutants gained 2.7-fold more mass than those feeding on wild-type plants (Fig. 4H). Consistently, acylsugar content in the frass of larvae fed on *abcb5* mutants was less than 0.4% of that in larvae fed on wild-type LA0716 (Fig. 4I). Collectively, these results demonstrate that SpABCB5 controls acylsugar secretion from trichome apical cells into both the external environment and the underlying mesophyll, thereby providing defense against leaf mining (endophagous) pests.

### Acylsugars confer resistance to free-living insects

To further test whether the acylsugar translocation procedure influences resistance against free-living (ectophagous) insect, we evaluated the performance of the common cutworm (*Spodoptera litura*), a highly polyphagous pest capable of feeding on over 100 host plant species including tomato [25]. Both the *asat1* biosynthetic mutant and the *abcb5* transport mutant exhibited significantly compromised resistance: larval feeding on these mutants gained approximately twofold more mass compared with larvae feeding on wild-type LA0716 plants (Fig. 5A and 5B). Complementation bioassays using an artificial diet supplemented with acylsugars at physiologically relevant concentrations confirmed that these metabolites significantly inhibit *S. litura* larval growth (Fig. 5C). Collectively, these results demonstrate that acylsugars are critical for conferring resistance against free-living insect herbivores.

**Fig. 5.**
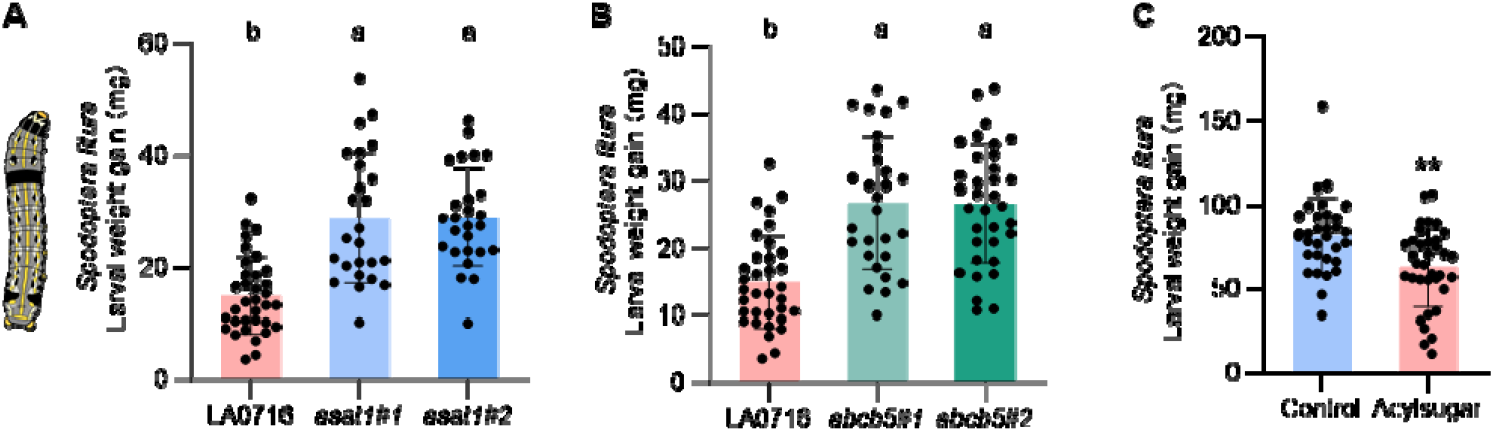
Acylsugars confer resistance against *Spodoptera litura* larvae. (A) Larval mass of *S. litura* (mean ± SE, *n* = 34, 27, and 28, respectively) after 10 days of feeding on LA0716 wild-type and *asat1* mutant plants. (B) Larval mass of *S. litura* (mean ± SE, *n* = 34, 28, and 34, respectively) after 10 days of feeding on LA0716 wild-type and *abcb5* mutant plants. (C) Larval mass of *S. litura* (mean ± SE, *n* = 32 and 35, respectively) after 3 days of feeding on artificial diet (Control) or artificial diet supplied with acylsugars (acylsugar). Different letters indicate significant differences (*P* < 0.05, one-way ANOVA with Tukey’s *post hoc* test). Statistical significance for C was determined using independent-sample student-*t* tests (\*\**P* < 0.01).

## Discussion

Trichomes serve as the first layer of mechanical and chemical barriers against free-living (ectophagous) insects [26-28]. In this study, we demonstrate that trichome-derived acylsugars, which are predominately secreted into the external environment, also provide robust defense against the endophagous tomato leafminer. Metabolite profiling of larval frass and mesophyll tissues via MALDI-MSI, combined with a promoter-reporter system driven by *ASAT1*, confirmed the translocation of trichome-synthesized acylsugars into the mesophyll tissues upon leafminer infestation. Furthermore, metabolite analysis and bioassays using *SpABCB5* knockout mutants identified *SpABCB5* as the key transporter mediating this trichome-to-mesophyll acylsugar translocation, a mechanism essential for the strong resistance observed in the acylsugar-rich wild tomato *Solanum pennellii* (Fig. 6). Collectively, our findings highlight the versatility of the trichome as a “metabolic biofactory” and underscore the critical contribution of epidermal defenses in protection against leaf mining pests.

**Fig. 6.**
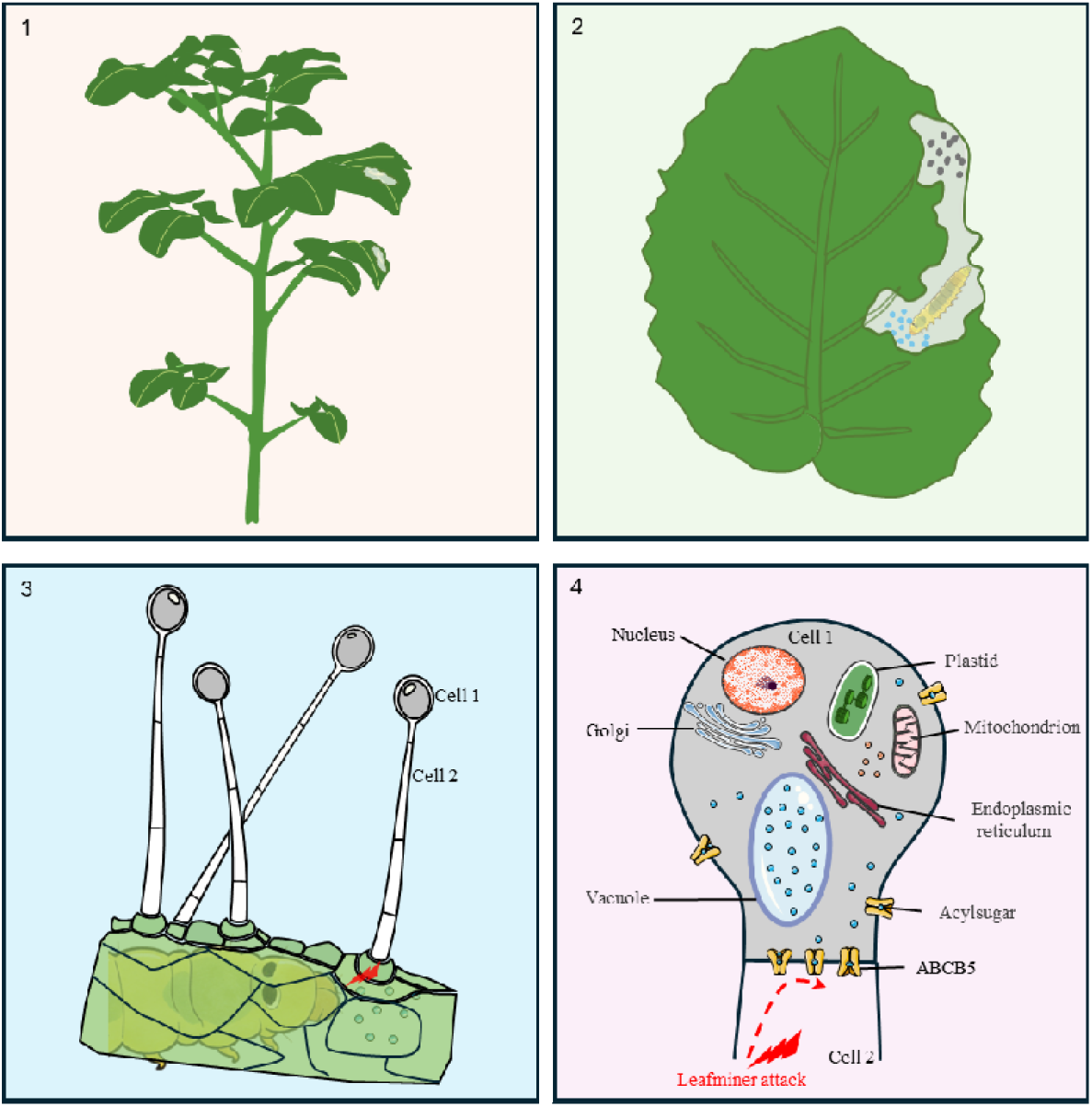
Model depicting the translocation of trichome-synthesized acylsugars into mesophyll tissues upon leafminer larval infestation.

ATP-binding cassette (ABC) transporters are integral membrane proteins that actively mediate the translocation of diverse substrates, including organic acids, metal ions, phytohormones and specialized metabolites [29]. Although hydrophobic compounds were historically thought to diffuse passively through lipid membranes [30], recent evidence indicates that plants require specific transporters for the secretion of such compounds. Notable examples include PhABCG1, which facilitates volatile emission from *Petunia* flowers [31] and AaPDR3, which mediates β-caryophyllene transport in *Artemisia annua* trichomes [32]. During the preparation of this manuscript, an independent study identified *SpABCB5* and its homolog in cultivated tomato can transport acylsugar *in vitro*, and mutant of which *in vivo* affects acylsugar accumulation in the external environment [33], consistent with our observation of the absence acylsugar droplets on the trichomes of the independent *abcb5* mutant lines. Our study demonstrates that ABCB5 mediates the efflux of acylsugar from the biosynthetic apical cells of trichomes, directing transport both apically toward the external droplet and basipetally toward the underlying mesophyll cells. Crucially, this basipetal translocation is induced by leafminer feeding, as acylsugar signals were virtually undetectable in the mesophyll via MALDI-MSI in unattacked plants. Thus, the bidirectional transport of acylsugars appears to be tightly regulated by a sophisticated polar transport mechanism. Polar transport is best characterized in plants in the context of the phytohormone, auxin [34]. The polar transport of auxin by PIN, the auxin efflux carriers, is a well-studied process governed by the polar localization of PIN proteins [35] and the modulation of their transport activity [36]. Concomitantly, we hypothesize that leafminer infestation triggers a signal transduction cascade from mesophyll into trichome apical cells, thereby re-organizing ABCB5 protein localization to the basipetal side of trichome apical cells or activating an unknown transport-activity rheostat. Previous studies showed that trichomes can act as sentinels, sensing mechanical stimuli such as insect crawling, that trigger Ca^2^□ waves. These mechanical signals are transmitted to trichome basal cells and subsequently to neighboring type-VI glandular trichome, initiating terpene biosynthesis via active jasmonic acid signaling [37]. A similar rapid metabolic response (within 6 h) in trichomes to *Manduca sexta* herbivory and jasmonate elicitation has been reported in *Nicotiana repanda*, in which the mesophyll nornicotine pool is transported to trichomes for the synthesis of the dramatically more toxic N-acyl-nornicotines that are deposited onto leaf surfaces after attack [38]. Future studies should elucidate the mechanisms underlying the unidirectional and inducible function of ABCB5 in acylsugar translocation.

Our stimulated Raman scattering (SRS) microscopy analysis revealed that acylsugars are also stored in vacuoles, indicating that while acylsugars predominantly accumulate as extracellular droplets on trichomes, a substantial fraction is retained in vacuoles. This evidence of vacuolar acylsugar storage challenges the prevailing assumption that acylsugars are secreted and remain on trichomes once synthesized [39]. SRS imaging is a powerful, gold-standard tool for elucidating the subcellular localization of plant specialized metabolites. Although the vacuole is typically considered as a dynamic repository of plant specialized metabolites [40], the majority of acylsugars are constitutively secreted and accumulated externally, contributing to physical defense against free-living pests such as *S. litura*, or as indirect defenses against *M. sexta* larvae by olfactorily tagging them for predation [41]. In this context, vacuolar storage is likely not a primary requirement. However, this storage pool may facilitate inducible translocation into the mesophyll upon leafminer infestation. We hypothesize that this translocated pool originates from vacuolar stores and/or *de novo* synthesis, rather than from apoplast droplets, which are less likely to rapidly respond to internal infestation signals. *S. pennellii* accumulates acylsugars up to 20% of leaf dry mass [42]. Our data show that leafminer frass contains acylsugars at approximately half the concentration found in leaves, a considerable amount that is unlikely produced solely by rapid *de novo* synthesis. Given that ABCB5 localizes to the plasma rather than tonoplast membrane (Fig. S12), and *abcb5* mutants accumulate increased vacuolar acylsugars, ABCB5 is likely not directly involved in vacuolar accumulation. Transporters from other families, such as the multidrug and toxic compound extrusion (MATE) transporter NtJAT1 from *Nicotiana tabacum* [43] and the nitrate/peptide family (NPF) transporter CrNPF2.9 from *Catharanthus roseus* [44], are localized to the tonoplast and facilitate the transport of nicotine and monoterpene indole alkaloid intermediate, respectively. Further research identifying the specific tonoplast-localized acylsugar transporter will clarify the ecological function of vacuole storage, particularly regarding defense against leaf mining pests.

The *abcb5* transporter mutant exhibited significantly reduced the resistance to free-living *S. litura*, a phenotype mirroring that of the *asat1* biosynthetic mutant. We attribute the susceptibility of the *abcb5* mutant to impaired acylsugar biosynthesis rather than to a defect in translocation, given that free-living caterpillars consume entire leaves including both trichomes and mesophyll tissues. In contrast, the compromised resistance of the *abcb5* mutant against the leafminer *P. absoluta* is influenced by its inability to translocate acylsugars from trichomes to mesophyll tissues. Specifically, while the *asat1* mutant accumulates only 0.2% of the acylsugar found in wild-type LA0716 leaves, frass from *P. absoluta* larvae fed on *asat1* contains approximately 5% of the acylsugar content found in the frass of larvae fed on wild-type plants. However, the *abcb5* mutant accumulates 5% of the wild-type acylsugar levels in leaves, yet frass from *P. absoluta* fed on *abcb5* contains only 0.4% of the frass of larvae fed on wild type plants. This inverse relationship between leaf and frass acylsugar levels indicates that SpABCB5 contributes to defense against leafminer not only by indirectly promote acylsugar biosynthesis, but also by directly by translocation trichome acylsugars into mesophyll tissues.

Given the devastating yield losses caused by the tomato leafminer, substantial efforts have been directed at identifying resistant tomato germplasm. Previous studies have established a correlation between pest resistance and acylsugar concentration [18-20, 45]. However, this resistance was largely explained as antixenosis (non-preference) rather than direct antibiosis. Similarly, trichomes conferred resistance against the leafminer fly, *Liriomyza trifolii*, another devastating invasive pest, was reported as a physical barrier to female oviposition, thereby causing pre-reproductive mortality [46]. The current study demonstrates the direct antibiosis of acylsugars against leafminer larval growth and elucidates the underlying mechanisms. Thus, these findings provide mechanisms for a trichome-derived metabolite-based management of endophagous pests. This work supports the utilization of acylsugars as environmentally friendly pesticides for broad-spectrum pest control and their application in crop metabolite engineering.

## Materials and Methods

### Plant material and growth conditions

Seeds of *Solanum lycopersicum* cv. M82 and *Solanum pennellii* accession LA0716 were obtained from the C.M. Rick Tomato Genetics Resource Center (University of California, Davis). The tomato and the genetically modified tomato lines were cultivated in a glasshouse under a photoperiod of 16-h light and 8-h darkness, with a temperature of 25°C. The *Nicotiana benthamiana* plants were cultivated under the same growth conditions.

Knockout mutants of *spasat1* and *spabcb5* in LA0716 were generated using CRISPR/Cas9 technology. Generation and genotyping of *spasat1* and *spabcb5* plants was performed according to method description in *SI Appendix, Materials and Methods*.

### VIGS

Vector construction, plant growth, and inoculation conditions for VIGS were performed following previously described procedures [47]. Detailed information is in *SI Appendix, Materials and Methods*.

### Imaging and counting of tomato trichomes by Super-Depth Digital Microscope

Fully expanded apical leaves from the four uppermost branches of 6-week-old tomato plants were sampled for trichome counting and imaging using DMS1000 3D Super-Depth Digital Microscope (NINGBO SUNNY, China). For type I/IV trichome counting, at least three fields on the adaxial surface of each leaf were imaged, and type I/IV trichomes in each image were counted. Counts from four leaves per seedling were averaged to obtain one biological replicate, and six biological replicates were analyzed.

### Acylsugars extraction and LC-MS analysis

The trichome acylsugars extracted from leaves were analyzed using Q Exactive quadrupole-orbitrap high-resolution mass spectrometry coupled with a Dionex Ultimate 3000 RSLC (HPG) ultra-performance liquid chromatography (UPLC-Q-Orbitrap-HRMS) system (Thermo Fisher Scientific), equipped with an electrospray ionization (ESI) probe operating in positive mode. Detailed acylsugars extraction and LC/MS methods are in *SI Appendix, Materials and Methods*.

### Acylsugars purification and bio-assay

Acylsugars used for compound feeding assays were isolated and purified from leaves of 8-week-old *S. pennellii* plants. *P. absoluta* used in bio-assay was originally collected from tomato fields in Xinjiang, China. Larvae were reared in a growth chamber at 25 °C under a 16-h light / 8-h dark photoperiod at CEMPS, Shanghai, and fed on *S. lycopersicum* cv. M82 plants. *Spodoptera litura* were purchased from Henan Jiyuan Baiyun Industrial Co., Ltd (China). Larvae were kept under the same growth conditions, and fed on artificial diet. The detailed steps for acylsugars purification and bio-assay are in *SI Appendix, Materials and Methods*.

### Plant treatment and gene expression analysis

Samples of roots, leaves, stems, leaf trichomes, stem trichomes from 5□week□old *S. pennellii* LA0716, as well as flowers at full bloom stage and 14 days post-anthesis fruits, were collected for RNA□seq to analyze tissue□specific expression pattern. Leaf trichomes of LA0716 and *spabcb5* mutants were collected for RNA□seq analysis to investigate the metabolic pathways in which *SpABCB5* is involved. The raw RNA-seq data have been deposited in the China National Genomics Data Center Genome Sequence Archive (accession number: PRJCA060947).

To screen transporters involved in the translocation of acylsugars into mesophyll, fully expanded apical leaves from the four uppermost branches of 4-week-old tomato plants were infested with third-instar *P. absoluta* larvae. After 24 hours, the leaves were harvested for quantitative real-time PCR (qRT-PCR) analysis. Details about RNA-seq, expression patterns analysis, and qRT-PCR are provided in the *SI Appendix, Materials and Methods*.

### Fluorescence Microscopy

Third-instar *P. absoluta* larvae were used to feed on mature leaves of 4-week-old *proSlASAT1::GFP* plants. GFP expression driven by the *SlASAT1* promoter was visualized using the laser scanning confocal microscope (Nikon, Japan). Fluorescence from GFP was detected by excitation at 488 nm and a 505-to 525-nm emission filter.

Vacuoles in glandular trichomes were observed and photographed under bright-field illumination using a laser scanning confocal microscope (Nikon, Japan), and vacuole size was quantified using ImageJ software.

For subcellular localization of SpABCB5 in *N. benthamiana* leaf epidermal cells, RFP and GFP were imaged using the laser scanning confocal microscope (Nikon, Japan). RFP fluorescence was observed after excitation using a 587-nm laser and detected using the 610-nm emission filter. Detailed Subcellular localization methods are in *SI Appendix, Materials and Methods*.

### Stimulated Raman scatter (SRS) microscopy

Trichomes from fully expanded apical leaves of the four uppermost branches of 6-week-old tomato plants were used for the *in-situ* visualization of acylsugars by stimulated Raman scattering (SRS) microscopy. The Raman signals of authentic standards including acylsugars, sugar (starch), oleic acid (OA), and bovine serum albumin (BSA) were compared to screen for the characteristic Raman peaks of acylsugars. SRS experiments were performed as described previously [48]. Detailed SRS methods are in SI Appendix, Materials and Methods.

### MALDI–MSI analysis

To visualize the spatial distribution of acylsugars in tomato leaf tissues in situ, we employed MALDI□MSI (matrix-assisted laser desorption/ionization mass spectrometry imaging). Third-instar *P. absoluta* larvae were inoculated onto mature leaves of 4-week-old LA0716 plants. After 24 h, the remaining leaves were used to MALDI–MSI analysis. The detailed steps for MALDI–MSI analysis are in *SI Appendix, Materials and Methods*.

### Statistical analysis

Data were analyzed using SPSS 20.0 (SPSS Inc, http://www-01.ibm.com/software/analytics/spss/). Unless otherwise stated, unpaired Student’s *t*-test was used for two groups of data analysis, and one-way ANOVA with post hoc Tukey HSD test was used for three or more groups of data analysis.

## ACKNOWLEDGMENTS

We are grateful to Prof. Pengxiang Fan for providing with *pASAT1::GFP* transgenic tomato line. This research was supported by the National Natural Science Foundation of China (W2412106, 32372635, 32302460), the China Postdoctoral Science Foundation (2022M723145) and the Science and Technology Commission of Shanghai Municipality (23ZR1481600).

## AUTHOR CONTRIBUTIONS

J.L. conceived the project; Z.S., X.P., Y.Z., D.M., Y.L., Y.F., Y.G., N.C., J.X. and W.Y. performed research; S.B., M.J., J.D and X.X. contributed to the methodology. J.L. and Z.S. analyzed the data; J.L., Z.S. and I.T.B. wrote the manuscript draft. All authors edited and commented on the manuscript.

## DECLARATION OF INTERESTS

The authors declare no competing interests.

## Notes

### Competing Interest Statement

The authors have declared no competing interest.

