## Supplementary file for "Trichome acylsugar defenses are equally effective against leaf-mining and free-living insects"

**This PDF file includes:**

Supplemental methods

Figures S1 to S12

Tables S1

SI References

**Other supporting materials for this manuscript include the following:**

Movies S1

**Supplemental methods**

**Plant material and growth conditions**

Seeds of *Solanum lycopersicum* cv. M82 and *Solanum pennellii* accession LA0716 were obtained from the C.M. Rick Tomato Genetics Resource Center (University of California, Davis). The tomato and the genetically modified tomato lines were cultivated in a glasshouse under a photoperiod of 16-h light and 8-h darkness, with a temperature of 25°C. The *Nicotiana benthamiana* plants were cultivated under the same growth conditions.

**Generation of CRISPR/Cas9-mediated gene knockout plants**

Knockout mutants of *asat1* and *abcb5* in LA0716 were generated using CRISPR/Cas9 technology. Target-specific sgRNAs were designed via the CRISPR-P v2.0 platform (http://crispr.hzau.edu.cn/CRISPR2/). The sgRNAs (*SpASAT1* sgRNA1: CCCTCGTACTTACATCGATAATC, sgRNA2: TTCTGAATCGCGCGCTACCATGG; *SpABCB5* sgRNA1: AAGGTAGTTGAACAAACGGTGGG, sgRNA2: GCCAGCGCTTGGTTGACAGTAGG) were introduced into the pHCE vector. Constructs were verified by Sanger sequencing and transformed into *Agrobacterium tumefaciens* strain GV3101 for plant genetic transformation.

**VIGS**

Total RNA was extracted from 100 mg leaves of *S. pennellii* leaves using the TransZol Up Plus RNA Kit (TransGen Biotech, China) according to the manufacturer’s instructions. Complementary DNA (cDNA) libraries were prepared using the HiScript III 1st Strand cDNA Synthesis Kit (+gDNA wiper) (Vazyme, China). Vector construction, plant growth, and inoculation conditions for VIGS were performed following previously described procedures [1]. Briefly, 300-bp fragments of *SpABCB5* (*Sopen03g001870*), *SpABCG6* (*Sopen04g005380*), and *SpABCG54* (*Sopen12g034820*) were designed using the VIGS tool (http://solgenomics.net/tools/vigs) to avoid off-target silencing. Specific fragments were amplified by polymerase chain reaction (PCR) using primer pairs listed in Table S1. Amplified fragments were cloned in the vector *pTRV2* to obtain *pTRV2-SpABCB5*, *pTRV2-SpABCG6*, and *pTRV2-SpABCG54*. *Agrobacterium tumefaciens* strain GV3101 was transformed by electroporation with the resulting plasmids. Seeds germinated for 48 hours were pressure-infiltrated with a mixture of *A. tumefaciens* containing *pTRV1* and either *pTRV2-SpABCB5, pTRV2-SpABCG6, pTRV2-SpABCG54*, *pTRV2-PDS* (Phytoene desaturase, was used as a positive control to monitor silencing progress) or *pTRV2* (Empty vector, as a control).

**Imaging and counting of tomato trichomes by Super-Depth Digital Microscope**

Fully expanded apical leaves from the four uppermost branches of 6-week-old tomato plants were sampled for trichome counting and imaging using DMS1000 3D Super-Depth Digital Microscope (NINGBO SUNNY, China). For type I/IV trichome counting, at least three fields on the adaxial surface of each leaf were imaged, and type I/IV trichomes in each image were counted. Counts from four leaves per seedling were averaged to obtain one biological replicate, and six biological replicates were analyzed.

**Acylsugars extraction and LC-MS analysis**

To analyze the relative contents of acylsugars in different plants, fresh tomato leaves were flash-frozen in liquid nitrogen and ground into a fine powder using a mortar and pestle. Approximately 100 mg of the powder was transferred into a 2 mL centrifuge tube, extracted with 10 volumes of 80% (v/v) aqueous methanol, and vortexed for 1 min. The mixture was centrifuged at 14,000×g and 4 °C for 30 min, and this procedure was repeated twice. Finally, the supernatant was transferred to a 2 mL chromatography autosampler vial and stored at −20 °C until LC‑MS analysis.

To determine the acylsugars content in leafminer frass, second-instar leafminer larvae reared on M82 plants were transferred to corresponding *S. pennellii* plants. After 48 hours of feeding, the larvae were removed and placed in centrifuge tubes for 12 h to allow defecation. The frass was collected, and acylsugars were extracted from the frass using the same method as for leaf acylsugars.

Metabolite analysis was conducted using Q Exactive quadrupole-orbitrap high-resolution mass spectrometry coupled with a Dionex Ultimate 3000 RSLC (HPG) ultra-performance liquid chromatography (UPLC-Q-Orbitrap-HRMS) system (Thermo Fisher Scientific), equipped with an electrospray ionization (ESI) probe operating in positive mode. The source and ion transfer parameters applied were as follows: spray voltage 3.5 kV. For the ionization mode, the sheath gas, aux gas, capillary temperature and heater temperature were maintained at 40, 10 (arbitrary units), 300 °C and 350 °C, respectively. The S-lens RF level was set at 50. The Orbitrap mass analyzer was operated at a resolving power of 70,000 in full-scan mode (scan range: 80-1200 m/z; automatic gain control (AGC) target: 1e6) and of 17,500 in the Top 6 data-dependent MS^2^ mode (stepped normalized collision energy: 20, 40 and 60; injection time: 50 ms; isolation window: 1.5 m/z; AGC target: 1e5) with a dynamic exclusion setting of 3.0 seconds. Chromatographic separation was performed on a ACQUITY UPLC HSS T3 column (100 mm × 2.1 mm, 1.8 μm particle size; Waters) column maintained at 40°C, with sample aliquots of 2 µL separated using a 30-minute binary gradient of 0.01% formic acid and 2 mM ammonium formate in water (mobile phase A) and methanol: acetonitrile (1:1, v/v, mobile phase B) at a flow rate of 0.3 mL/min. The gradient elution programs were set as follows: 0.0-1.0 min, 2.0% B; 1.0-25.0 min, 2.0%-98.0% B; 25.0-27.0 min, 98% B; 27.1-30.0 min, 2.0% B.

**Acylsugars purification for bio-assays**

Acylsugars used for compound feeding assays were isolated and purified from *Solanum pennellii* as follows. 100 g leaves from 8-week-old plants were harvested and extracted using a dipping buffer consisting of methanol:water (8:2, v/v). Plant tissues were submerged in 200 mL of the dipping buffer for 1 min with gentle agitation to ensure thorough extraction. The solvent was concentrated to less than 50 mL under reduced pressure using a rotary evaporator, and the concentrate was centrifuged at 10,000×g for 30 min twice to obtain the crude extract. Crude extract separation was performed using an Agilent 1260 Infinity semi-preparative HPLC system. Chromatography was conducted on a Welch Ultimate XB-C18 column (21.2 mm × 250 mm, 5 µm particle size) maintained at 40 °C. The sample injection volume was 5 mL. The mobile phase consisted of (A) 0.1% (v/v) formic acid in water and (B) acetonitrile. A linear gradient was applied as follows: 5% B (0–5 min), increasing to 95% B (5-55 min), increasing to 100% B (55-67 min), then to 5%B (67-69min). The column was re-equilibrated at 5% B from 69 min to 85 min. The flow rate was 10 mL/min. Eluted fractions were collected automatically at 1-minute intervals. All collected fractions were analyzed by LC-MS to assess purity and relative abundance of acylsugars. The fraction containing the purified acylsugars were concentrated to dryness under reduced pressure using a rotary evaporator and stored at -20°C until further use in compound feeding assays.

***Phthorimaea absoluta* and *Spodoptera litura* growth conditions and larval performance assays**

*P. absoluta* was originally collected from tomato fields in Xinjiang, China. Larvae were reared in a growth chamber at 25 °C under a 16-h light / 8-h dark photoperiod at CEMPS, Shanghai, and fed on *S. lycopersicum* cv. M82 plants. *S. litura* were purchased from Henan Jiyuan Baiyun Industrial Co., Ltd (China). Larvae were kept under the same growth conditions, and fed artificial diet.

To measure the performance of *P. absoluta* on plants (M82, LA0716, *pTRV2*, *pTRV2-ABCB5*, *pTRV2-ABCG6*, *pTRV2-ABCG54*, *asat1*, and *abcb5*), newly hatched neonates were reared on the indicated plants, and larval mass was measured on the 10^th^ day. Representative larval images were captured on the 10^th^ day.

To measure the performance of *S. litura* on plants (M82, LA0716, *asat1*, and *abcb5*), second–third instar larvae (6±2 mg) were reared on tomato seedlings and then collected and weighed on day 4. For compound feeding assays, size-matched second–third instar larvae were reared on the prepared artificial diet with or without 0.76 mg·g⁻¹ acylsugars, larval weights were recorded after 4 days. The artificial diet contained the following components per kilogram: soybean flour (100 g), wheat germ (80 g), yeast powder (26 g), casein (8 g), vitamin C (8 g), choline chloride (1 g), sorbic acid (2 g), cholesterol (0.2 g), inositol (0.2 g), and agar (26 g).

**Plant** **treatment and gene expression analysis**

Samples of roots, leaves, stems, leaf trichomes, stem trichomes from 5‑week‑old *Solanum pennellii* LA0716, as well as flowers at full bloom stage and 14 days post-anthesis fruits, were collected for RNA‑seq to analyze tissue‑specific expression pattern. Leaf trichomes of LA0716 and *spabcb5* mutants were collected for RNA‑seq analysis to investigate the metabolic pathways in which *SpABCB5* is involved. The raw RNA-seq data is deposited in the China National Genomics Data Center Genome Sequence Archive (accession number: PRJCA060947). A total of 31,637 expressed transcripts in various tissues were clustered into 20 clusters according to their expression patterns using Mfuzz, which is an improved algorithm based on Fuzzy C-Means. Gene expression profiles were analyzed and visualized as heatmaps using TBtools-II [2].

To screen transporters involved in the translocation of acylsugars into mesophyll tissues, fully expanded apical leaves from the four uppermost branches of 4-week-old tomato plants were infested with third-instar *P. absoluta* larvae. After 24 h, leaves were harvested for quantitative real-time PCR (qRT-PCR) analysis. qRT-PCR was performed using SYBR®Green Realtime PCR Master Mix (TOYOBO, Japan) on a CFX96™ real-time system. The *SpActin* was used as an internal control. Four biological replicates were analyzed for each sample. Primers used for expression analysis of target genes are listed in Table S1.

**Fluorescence Microscopy**

Third-instar *P. absoluta* larvae were used to feed on mature leaves of 4-week-old *proSlASAT1::GFP* plants. GFP expression driven by the *SlASAT1* promoter was visualized using the laser scanning confocal microscope (Nikon, Japan). Fluorescence from GFP was detected by excitation at 488 nm and a 505- to 525-nm emission filter.

Vacuoles in glandular trichomes were observed and photographed under bright-field illumination using a laser scanning confocal microscope (Nikon, Japan), and vacuole size was quantified using ImageJ software.

**Subcellular localization**

To analyze the subcellular localization of SpABCB5, the coding sequence (CDS) fragment of *SpABCB5* without the stop codon was amplified by PCR (primers used are listed in Supplementary Table) and then inserted into the *pCAMBIA1300-GFP* vector to produce the fusion construct *pCAMBIA1300-SpABCB5-GFP* using the ClonExpress Ultra One Step Cloning Kit V2 (Vazyme, China). Then, *pCAMBIA1300-SpABCB5-GFP* and PM-RFP marker based on *AtPIP2A* were transformed into *A. tumefaciens* strain GV3101 and injected into 4-week-old *Nicotiana benthamiana* leaves [3]. The green fluorescence of GFP and the red fluorescence of RFP were observed and captured by a laser confocal microscope (Nikon, Japan) at 48 h post-infiltration. The fluorescence signal intensity was calculated using ImageJ software.

**Stimulated** **Raman scatter (SRS) microscopy**

Trichomes from fully expanded apical leaves of the four uppermost branches of 6-week-old tomato plants were used for the *in-situ* visualization of acylsugars by stimulated Raman scattering (SRS) microscopy. The Raman signals of authentic standards including acylsugars, sugar (starch), oleic acid (OA), and bovine serum albumin (BSA) were compared to screen for the characteristic Raman peaks of acylsugars.

SRS experiments were performed as described previously [4]. Spontaneous Raman scattering spectra were acquired including a monochromator (iHR320, Horiba), a charge-coupled device camera (Symphony, Horiba) and a microscope (IX71, Olympus) with a 40 × air objective with 633 nm helium-neon laser beam at room temperature. Raman spectra were acquired at a single point for all samples under identical conditions, using a 10 s integration time and averaging 10 scans per spectrum. Quartz plate was used to suppress autofluorescence generated by glass slides. For the SRS microscope, a commercial femtosecond laser system (Insight DS+, Spectra-Physics) produced two synchronized pulse trains at 80 MHz. The fixed fundamental output of 1040 nm was employed as the Stokes beam (~ 200 fs), while the tunable optical parametric oscillator output (680 to 1300 nm, ~ 150 fs) served as the pump beam. To acquire high spectral resolution (~ 13 cm-1), pulse durations of the pump and Stokes beams were chirped and stretched by passing through SF57 glass rods (~ 3.8 ps for the pump pulse and ~ 1.8 ps for the Stokes). The intensity of the stokes beam was modulated at 1/4 of the laser pulse repetition rate (80 MHz) using a polarizing beam splitter (PBS) and an electrooptical modulator (EOM, EO-AM-R-20-C2, Thorlabs). The two laser beams were spatially and temporally overlapped via a dichroic mirror, then delivered into a laser scanning microscope (FV1200, Olympus) equipped with galvo mirrors for raster scanning. The combined beams were focused onto the sample by a 60 × water immersion objective lens (Olympus, UPLSAPO 60XWIR, NA 1.2). The transmission of the forward-going pump and Stokes beams was collected by a high N.A. oil condenser (oil immersion, NA=1.4, Nikon) after passing through the sample. Transmitted through a bandpass filter (CARS ET890/220, Chroma), the stimulated Raman loss (SRL) signal was detected by a homemade reverse-biased photodiode (PD) and demodulated with a lock-in amplifier (HF2LI, Zurich Instruments) at 20MHz to feed the analog input of the microscope to form images. The Raman band of C-H bonds was recorded at the pump laser of 802 nm, while the characteristic Raman peaks of acylsugars (1572-1615 cm^-1^) was measured with the tuned 896 nm pump beam. All the images used the same setting of 512 × 512 pixels with a pixel dwell time of 2 μs. The lateral resolution of the microscope system is ~350 nm, and the depth resolution is ~1 µm. Laser powers at the sample were: pump 30 mW and Stokes 40 mW.

**MALDI–MSI analysis**

To visualize the spatial distribution of acylsugars in tomato leaf tissues *in situ*, we employed MALDI‑mass spectrometry imaging. Third-instar *P. absoluta* larvae were inoculated into mature leaves of 4-week-old LA0716 plants. After 24 h, the remaining leaf tissues were punched into leaf discs. The lower epidermis was removed from half of each disc to expose the mesophyll, and the discs were then mounted upside down onto glass slides using conductive double-sided tape. A two-step matrix application method was used for α-cyano-4-hydroxycinnamic acid (CHCA): matrix sublimation was first performed using an iMLayer sublimation system (conditions: CHCA, 220 °C, 0.7 µm), followed by spraying 800 µL of 1 mg/mL CHCA matrix solution onto the slides with an airbrush. The slides were then air-dried at room temperature, and subjected to MALDI-MSI (matrix-assisted laser desorption/ionization mass spectrometry imaging) analysis.

All MALDI–MSI data were acquired utilizing an iMScope QT instrument (Shimadzu, Kyoto, Japan) equipped with an integrated optical microscope (magnification: ×5, ×10, and ×40), an atmospheric-pressure MALDI source, and a quadrupole-time-of-flight analyzer. The scanning area was determined using the optical microscope, and the sample sections were irradiated with a 355-nm Nd:YAG laser, employing the following parameters: laser shots, 700; repetition rate, 2000 Hz; laser diameter, 0 (5 μm); laser intensity, 45; detector voltage, 2.38 kV; and pitch (spatial resolution), 30 × 30 μm. The instrument was operated in positive mode, and spectra were acquired within the m/z range 300-800. The laser was calibrated using ink, ensuring accurate mass calibration of the MALDI–MSI instrument with the CHCA matrix prior to each experiment. All acquired data were subjected to analysis using IMAGEREVEAL MS software (Shimadzu, Kyoto, Japan). The same software was employed for data visualization and relative quantitative analysis. A tolerance of 10 ppm was set for MALDI–MSI, allowing exclusion of ions adjacent to the target ion to minimize interference.

**Multiple sequence alignment**

Protein sequences of SpASAT1 or SpABCB5 and corresponding mutants were aligned using MUSCLE (<https://www.ebi.ac.uk/jdispatcher/msa/muscle5>).

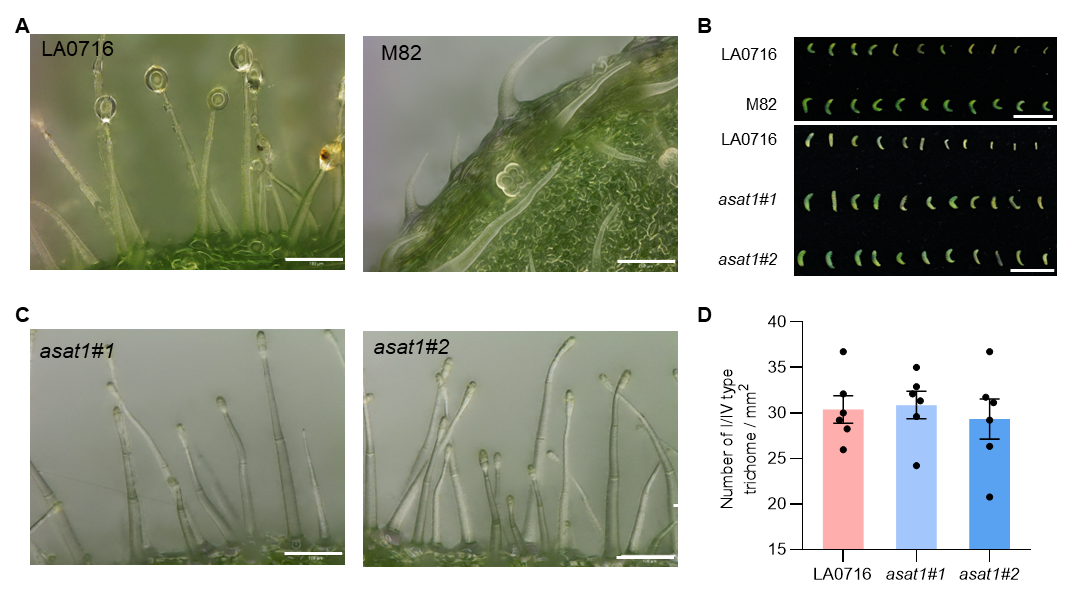

**Figure S1. Trichome morphology and *P. absoluta* larval performance across tomato genotypes, related to Figure 1.**

(A) Representative images of trichomes on leaves of wild tomato *S. pennellii* LA0716 and cultivated tomato M82. Scale bare, 100 μm. (B) Representative images of *P. absoluta* fed on leaves of different tomato genotypes. Scale bar, 1 cm. Weights are presented in Figs 1E and 2E. (C) Representative images of trichomes on leaves of independent *asat1* mutant lines. Scale bar, 100 μm. (D) Trichome densities (mean ± SE, *n* = 6) on indicated genotypes. No significant difference was detected using one-way ANOVA with Tukey’s *post hoc* test (*P* > 0.05).

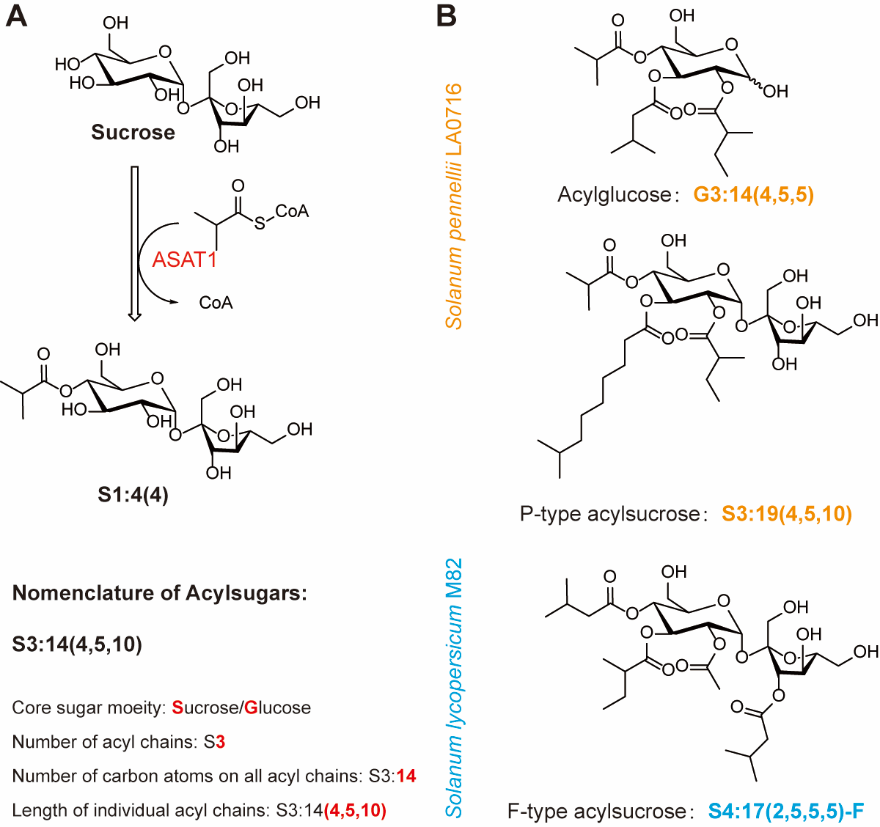

**Figure S2. Chemical structures and nomenclature of acylsugars, related to Figure 1.**
(A) Schematic illustrating that ASAT1 mediates the first step of acylsugar biosynthesis by transferring an acyl moiety onto the sucrose core. The nomenclature system for acylsugars is shown. (B) Structures of typical acylsugars found in *S. pennellii* (highlighted in orange) and *S. lycopersicum* (highlighted in blue). Acyl-sucroses harboring acyl moieties exclusively on the pyranose ring are defined as P-type acyl-sucrose. Those acylated on the furanose ring are defined as F-type acyl-sucroses.

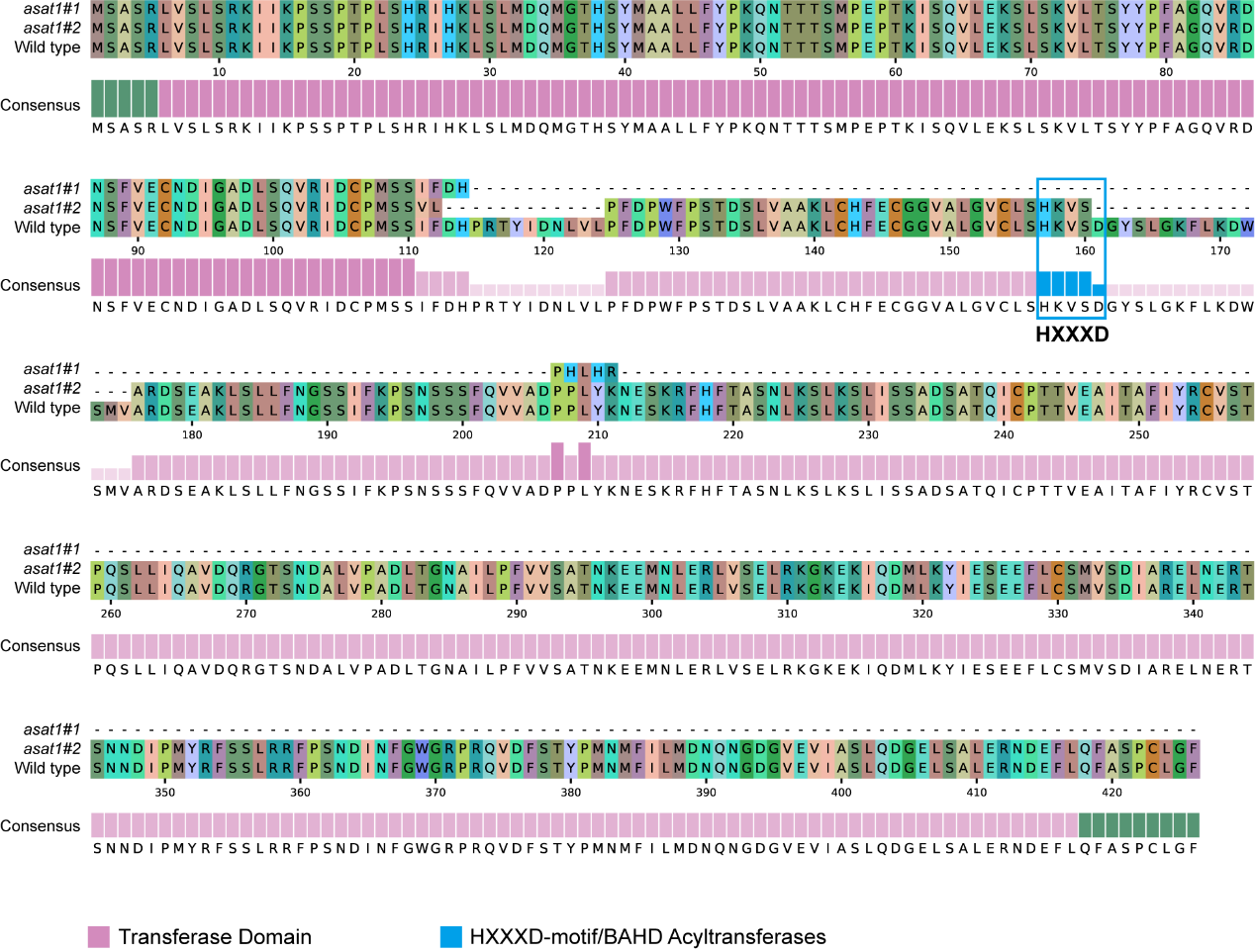

**Figure S3. Protein alignment of SpASAT1 and the *asat1* mutant sequence, related to Figure 2.**

The transferase domain is highlighted in pink, and the conserved HXXXD motif is highlighted in blue.

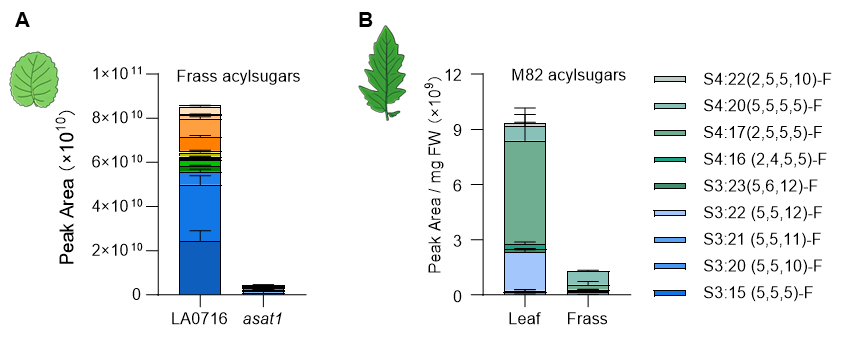

**Figure S4. Relative abundance of acylsugars in leaf extracts and larval frass, related to Figure 2.**
(A) Relative abundance of acylsugars (mean + SE, *n* = 3) in extracts of frass from larvae fed on wild-type LA0716 and *asat1* mutant plants. (B) Relative abundance of acylsugars (mean + SE, *n* = 3) in extracts of M82 leaves and frass from larvae fed on M82 plants.

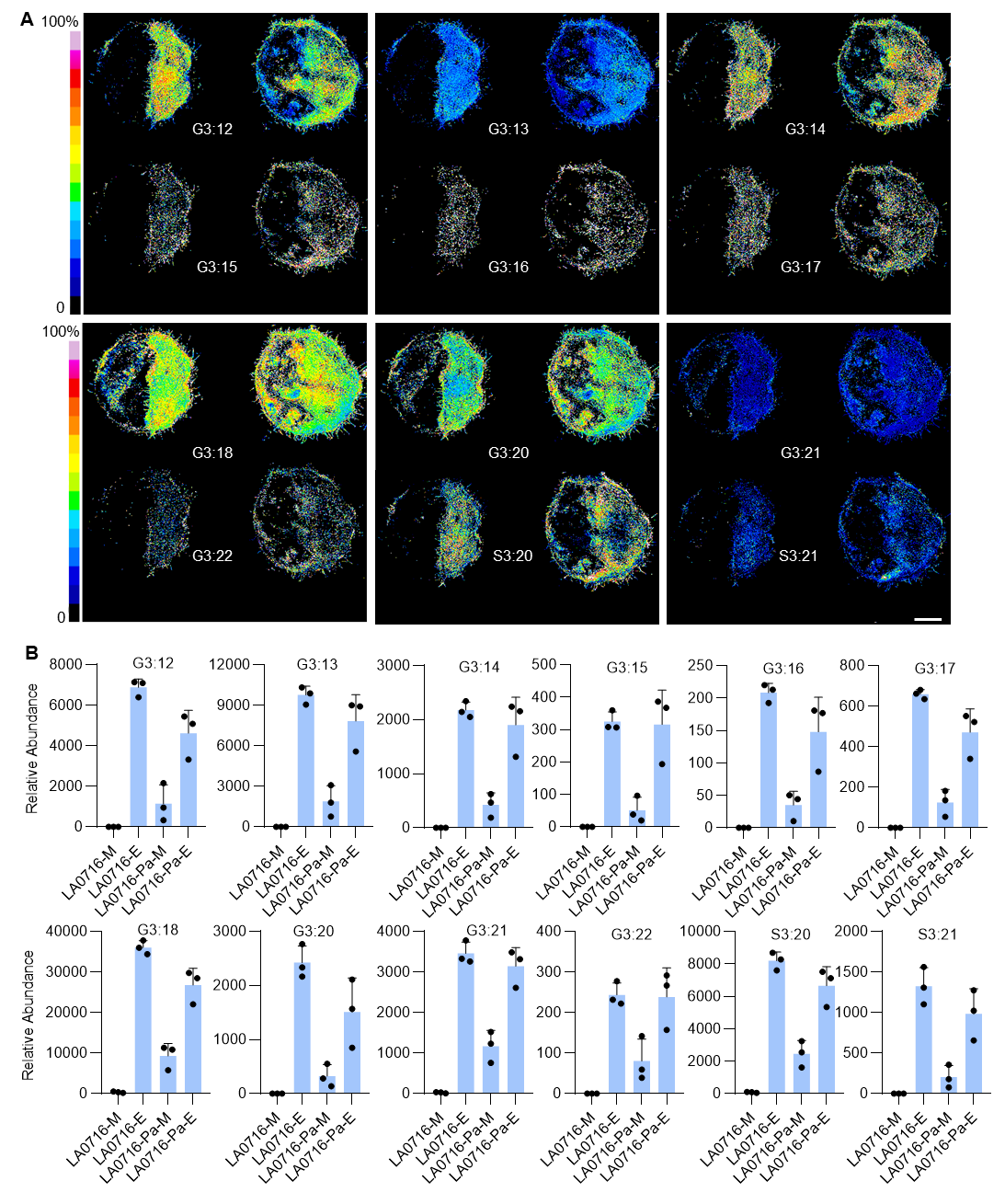

**Figure S5. Spatial distribution of acylsugars in the *S. pennellii* leaf in response to *P. absoluta* infestation, related to Figure 3.**

(A) MALDI-MSI of indicated acylsugars. The intensity of spectra representing each metabolite is visualized in false color. For each acylsugar, the left disc represents an uninfested control leaf, and the right disc represents a *P. absoluta*-infested leaf. For each leaf disc, the left half indicates MALDI-MSI signals harvested from the mesophyll (epidermis pealed), and the right half indicates signals harvested from the epidermis. Scale bar, 2 mm. (B) Relative intensity of MALDI-MSI signals (mean + SE, *n* = 3) for indicated acylsugars from indicated treatments. LA0716-M, mesophyll of uninfested LA0716; LA0716-E, epidermis of uninfested LA0716; LA0716-Pa-M, mesophyll of *P. absoluta*-infested LA0716; LA0716-Pa-E, epidermis of *P. absoluta*-infested LA0716.

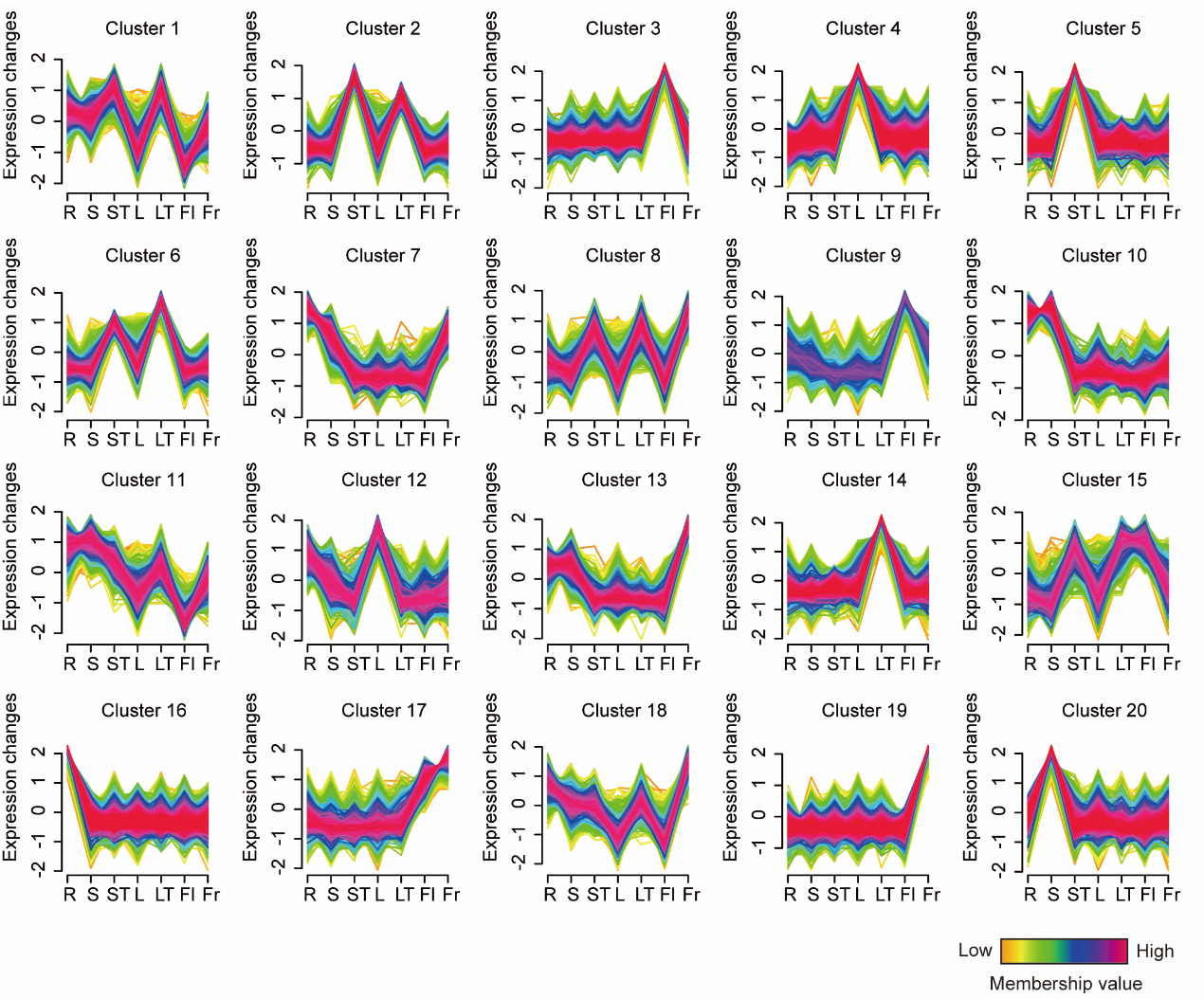

**Figure S6. Clustering analysis of transcriptome data from different LA0716 tissues, related to Figure 4.**

K-means clustering analysis divided the expressed genes into 20 clusters based on their expression patterns across different tissues. Cluster 2 represents the "trichome-specific" expression pattern. L, leaf; LT, leaf trichome; R, root; S, stem; ST, stem trichome; Fl, flower; Fr, fruit.

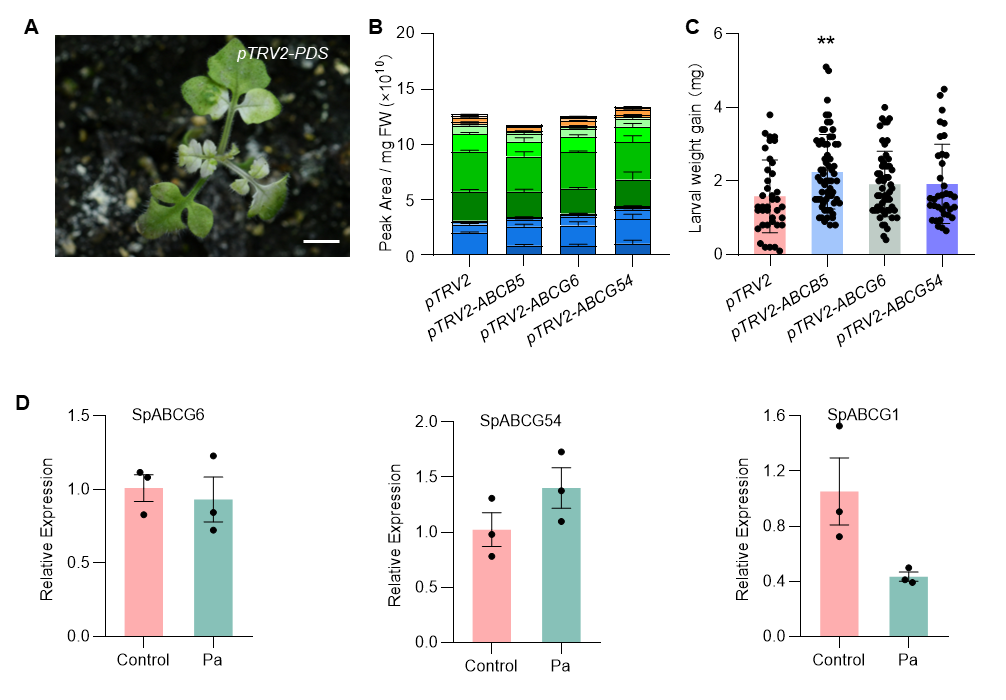

**Figure S7. Silencing candidate ABC transporter genes via VIGS, related to Figure 4.**

(A) Representative images of LA0716 plants with *PDS* gene silenced via VIGS (showing photobleaching). (B) Relative abundance of acylsugars (mean + SE, *n* = 3) in leaf extracts from indicated VIGS plants. (C) Larval mass (mean ± SE, *n* = 39, 63, 56, and 36, respectively) of *P. absoluta* after feeding on indicated plant genotypes for 10 days. Statistical significance for C was determined using independent-sample student-*t* tests (***P* < 0.01).

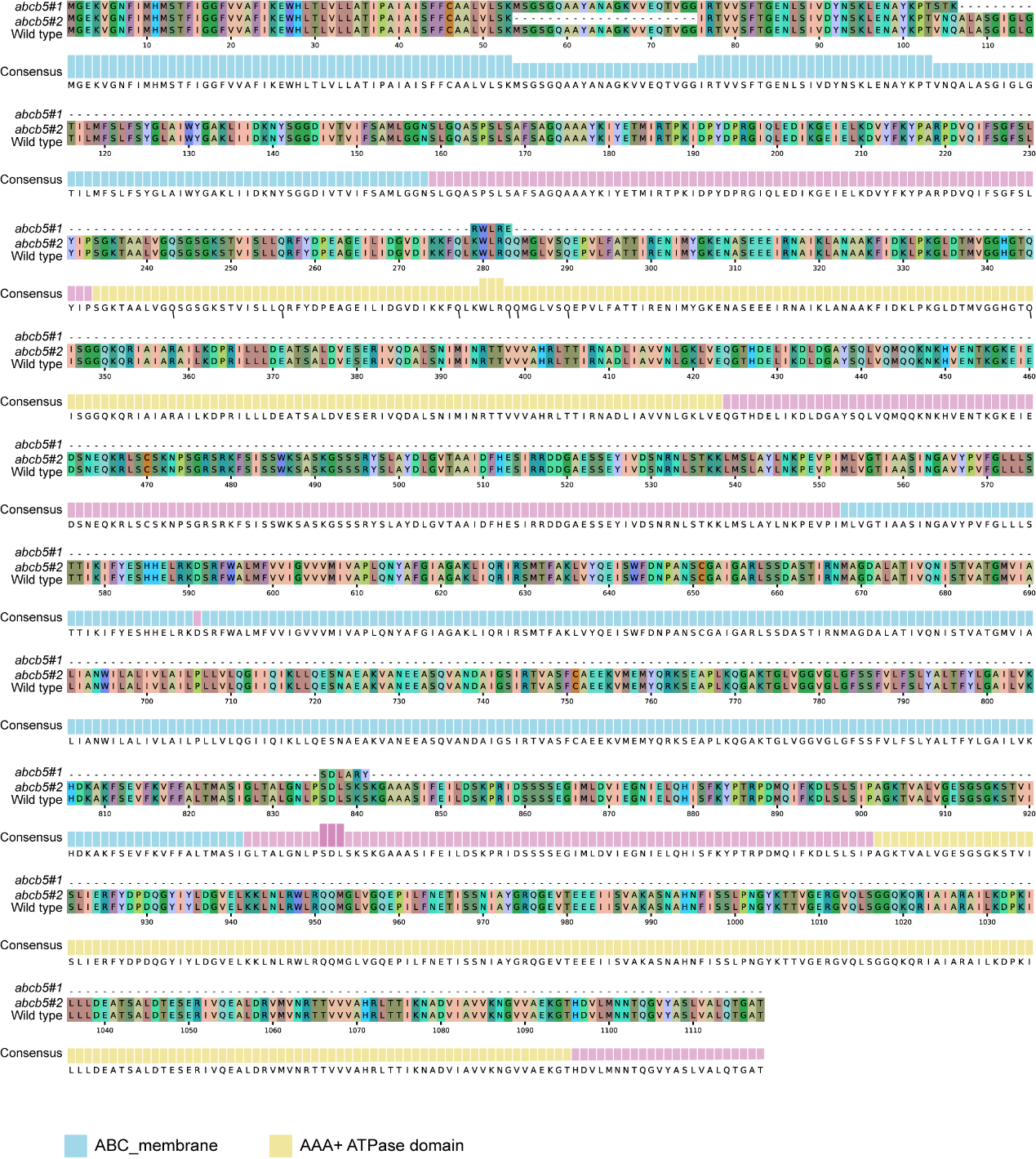

**Figure S8. Protein alignment of SpABCB5 and the *abcb5* mutant sequence, related to Figure 4.**

Different protein domains are indicated by different colors.

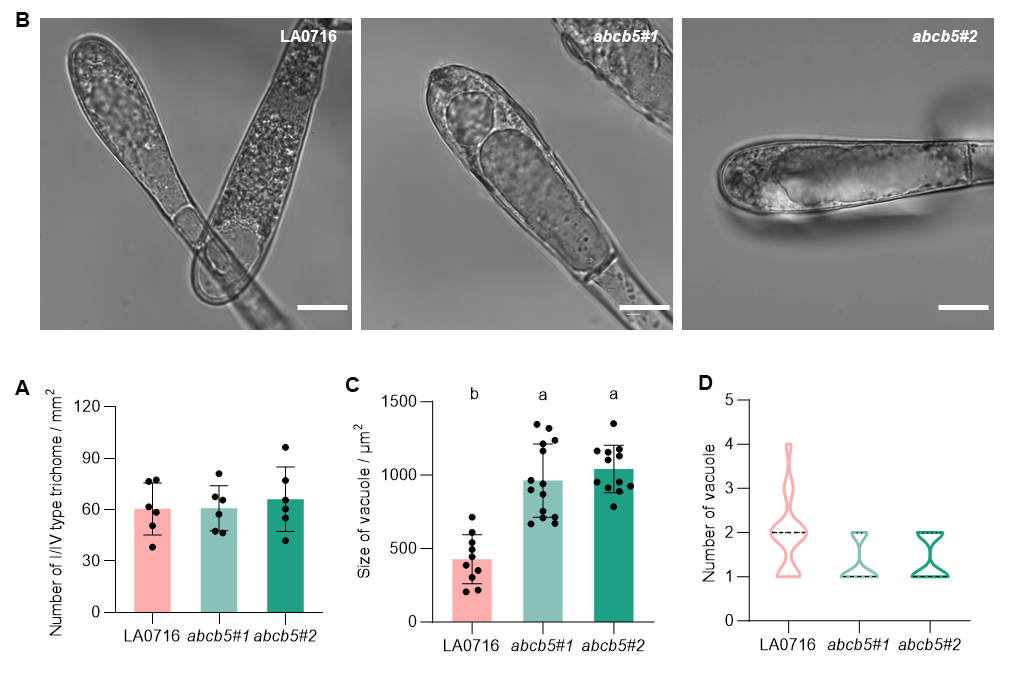

**Figure S9. Phenotypic characterization of the *abcb5* mutant, related to Figure 4.**

(A) Trichome densities (mean ± SE, *n* = 6) on indicated genotypes. (B) Microscopic image of trichome apical cells of indicated genotypes. Scare bar, 20 μm. (C) Vacuole size (mean ± SE, *n* = 11, 12 and 15, respectively) in the trichome apical cells of indicated genotypes. (D) Number of vacuoles (mean ± SE, *n* = 18, 18 and 19, respectively) per trichome apical cell. Different letters indicate significant differences (*P* < 0.05, one-way ANOVA with Tukey’s *post hoc* test).

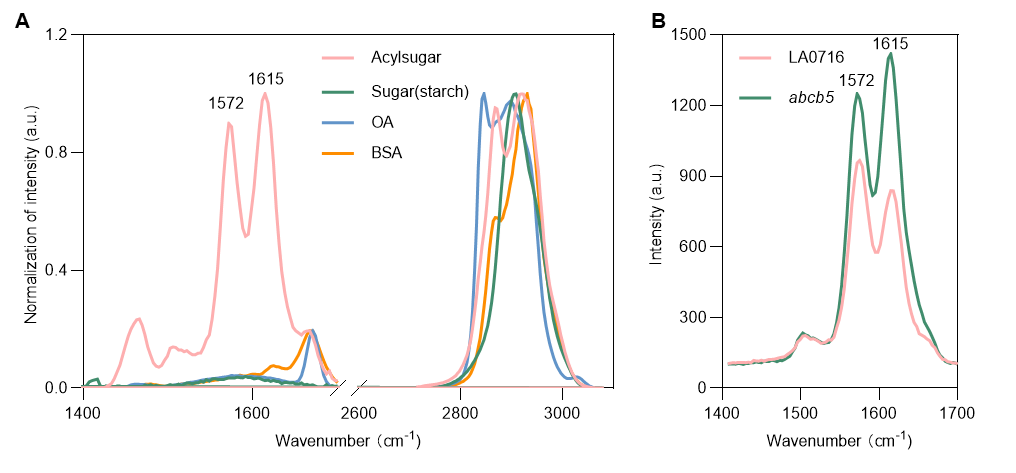

**Figure S10.** **SRS microscopy assay for acylsugar detection, related to Figure 4.**

(A) SRS spectra of pure acylsugar standard, starch, oleic acid (OA), and bovine serum albumin (BSA), showing distinct vibrational peaks at 1572 cm⁻¹ and 1615 cm⁻¹ specific to acylsugars. (B) SRS images of trichome apical cells from wild-type LA0716 and *abcb5* mutants.

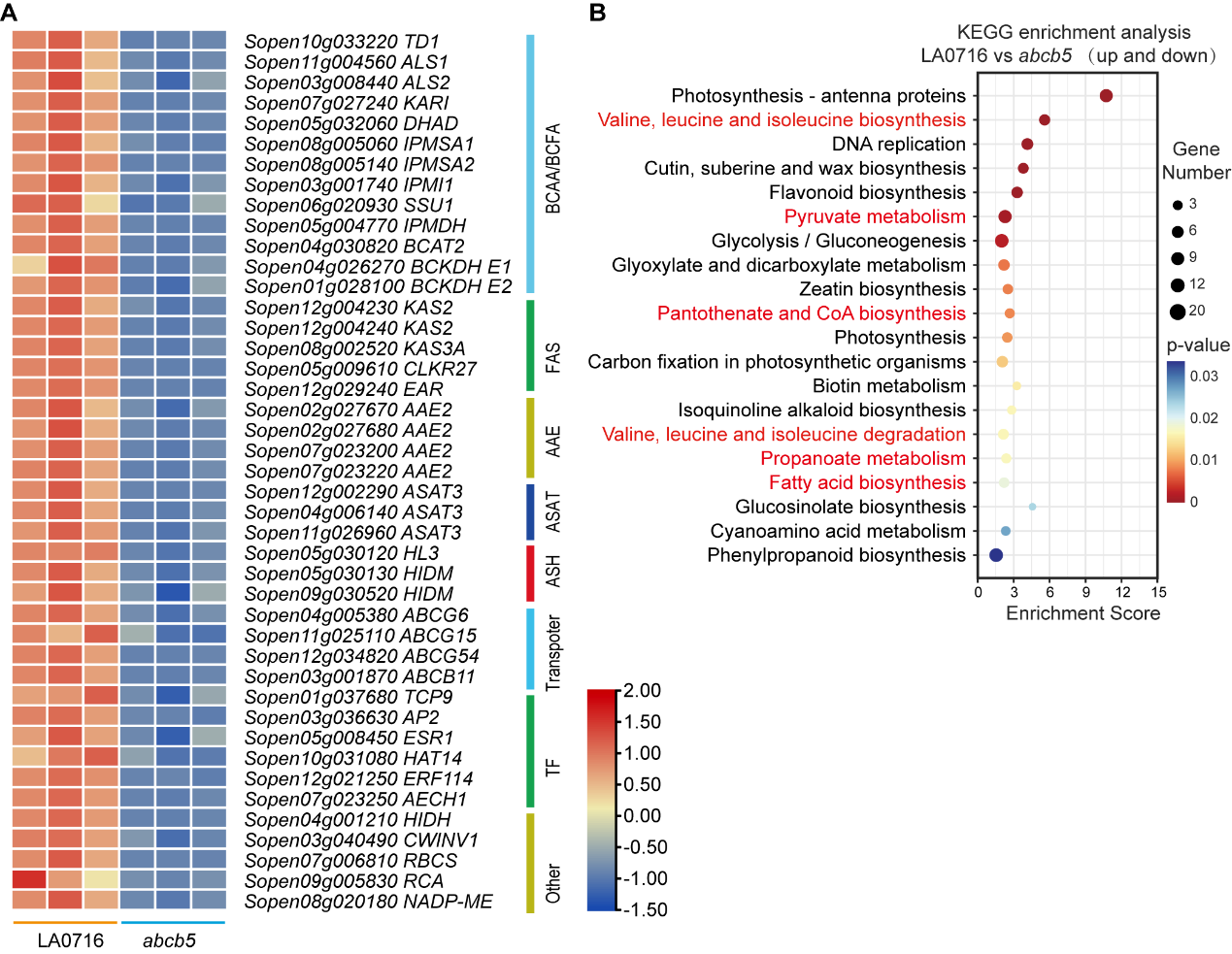

**Figure S11. Transcriptome analysis of *abcb5* mutants, related to Figure 4.**

(A) Heatmap of differentially expressed genes (DEGs) involved in acylsugar metabolism pathways in *abcb5* mutants compared to wild-type LA0716. (B) KEGG enrichment analysis of DEGs. Enriched metabolic pathways related to acylsugar biosynthesis are highlighted in red.

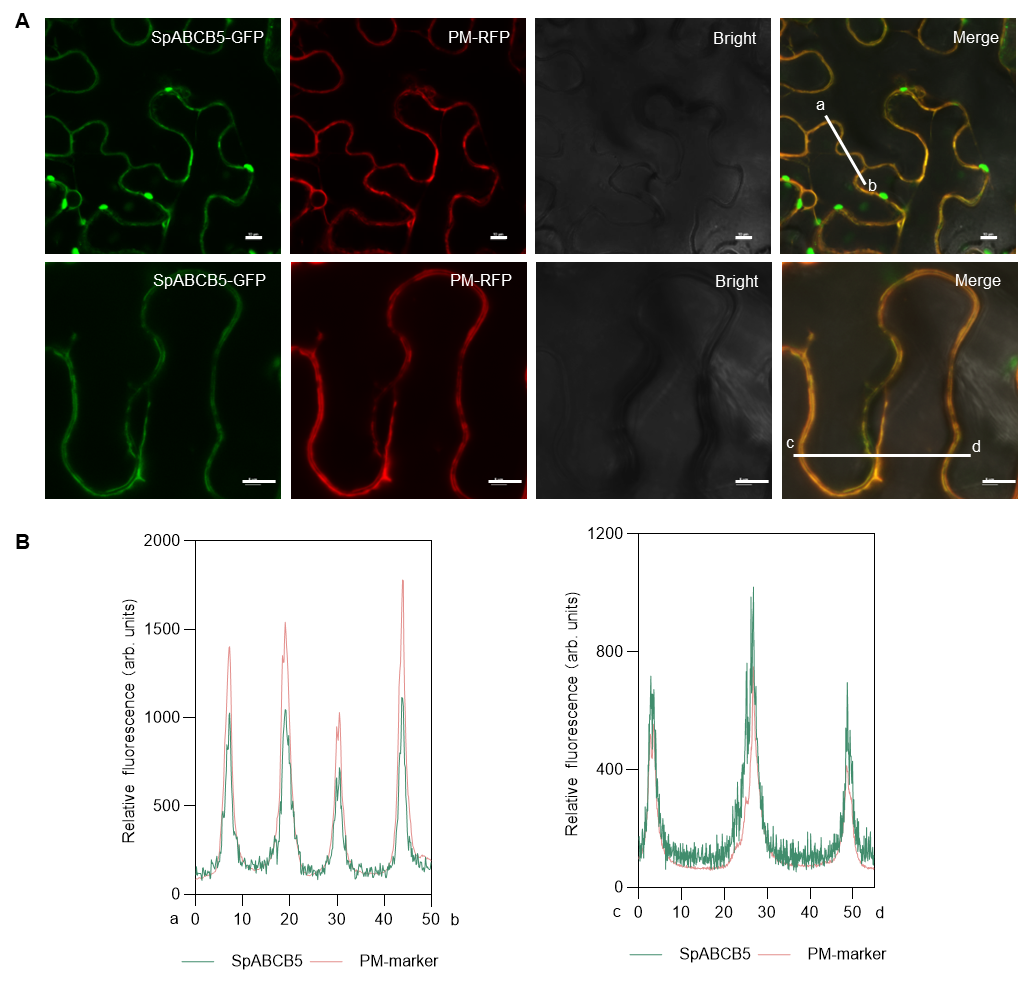

**Figure S12. SpABCB5 is localized at the plasma membrane,** **related to Figure 4.**

(A) SpABCB5-GFP colocalized with plasma membrane (PM) marker, *AtPIP2A*, which is labelled with RFP fluorescence. (B) Fluorescence intensity (in arbitrary units, arb. units) in cross-section [dotted line in (A)]. The turquoise lines indicate the intensities of SpABCB5-GFP, and the pink lines indicate the intensities of PM-RFP.

**Supplemental Table 1. Primers used in this study**

| **Primer name** | **Sequence (5’ - 3’)** | **Purpose** |
| --- | --- | --- |
| pTRV2-SpABCB5-F | gtgagtaaggttaccgaattcATAAGCATTTTCCAGCTTACTGTTATAA | Ligate fragments into pTRV2 |
| pTRV2-SpABCB5-R | cgtgagctcggtaccggatccATGGGTGAGAAGGTTGGGAATT |  |
| pTRV2-SpABCG6-F | gtgagtaaggttaccgaattcCGCGAGTAGTTCTCCGTCACG |  |
| pTRV2-SpABCG6-R | gggacatgcccgggcctcgagGATAATGTCACCGGCGACGA |  |
| pTRV2-SpABCG54-F | gtgagtaaggttaccgaattcTTTACTAATATGAGTAGCAGGAAATGTTTG |  |
| pTRV2-SpABCG54-R | gggacatgcccgggcctcgagCCATGATACTATAGAAGATGGAGGAAA |  |
| SpABCB11-GFP-F | acgggggactctagaggatccATGACACTTATTTTTGGTGACCTTGT | Ligate CDS into pCAMBIA1300-GFP |
| SpABCB11-GFP-R | gcccttgctcaccatggatccAGTTGCTCCTGTTTGAAGTGCA |  |
| qPCR-SpABCB11-F | GGTTCGACAACCCTGCAAAC | quantitative RT-PCR |
| qPCR-SpABCB11-R | CGCCAGCACTATAAGAGCGA |  |
| qPCR-SpActin-F | CCAGGCTGTGCTTTCTCTGT |  |
| qPCR-SpActin-R | CACGACCAGCAAGGTCCAAA |  |
| qPCR-SpABCG6-F | TTGCCGTTGGTACCCTCTTC |  |
| qPCR-SpABCG6-R | TAGCCGGTGACAAATCCACC |  |
| qPCR-SpABCG54-F | CATGCCTTGCCCTGTCAGAT |  |
| qPCR-SpABCG54-R | AAAGGCGTAAGGGATGGCAG |  |
| ASAT1-T1-T2-F | GCATGCCAGAACCAACGAAG | Genotyping mutant |
| ASAT1-T1-T2-R | TCCGCGTTGATCTACTGCTT |  |
| ABCB11-T1-F | ggagtgagtacggtgtgcCAACCATTCCTGCAATCGCC |  |
| ABCB11-T1-R | gagttggatgctggatggTGAGAGAACAAAGTACAATTCACCA |  |
| ABCB11-T2-F | ggagtgagtacggtgtgcTCTTCCTTTCCCTAGGTTGTTTC |  |
| ABCB11-T2-R | gagttggatgctggatggACAATGTCACCTCCGCTGTA |  |
| ASAT1-T1-T2-F | GCATGCCAGAACCAACGAAG |  |
